# Protist quantitative stable isotope probing identifies diverse active grazers in natural freshwater communities

**DOI:** 10.64898/2026.04.01.713104

**Authors:** Sofia Papadopoulou, Javier Florenza, Christoffer Bergvall, Eva S. Lindström, William D. Orsi

## Abstract

Bacterivorous protists are central to aquatic food webs, mediating the transfer of carbon and nutrients to higher trophic levels through the microbial loop. In natural communities, a major challenge remains in linking protist grazing activity to environmental sequences and identifying which taxa are actively feeding at the community level. Here, we present the first application of quantitative stable isotope probing (qSIP) in a grazing experiment. By combining qSIP with 18S rRNA gene amplicon sequencing, we linked prey assimilation to the identity of active protist predators at the operational taxonomic unit (OTU) level. In a replicated 36-h bottle-experiment, live ^13^C, ^15^N-labeled *Limnohabitans planktonicus* cells were added to natural samples from a lake pelagic site and its main inlet stream. Although hydrologically connected, protist richness was higher in the inlet than in the lake, yet a similar number of taxa incorporated prey biomass, comprising 108 OTUs in the inlet and 107 OTUs in the lake, including both rare and abundant taxa. Of these, 26 OTUs were labeled at both sites. The most strongly labeled protist in the inlet was a putative phago-mixotrophic prasinophyte, whereas in the lake it was an uncultured chrysophyte. Across sites, prey incorporation occurred in a broad range of taxa, including heterotrophs (e.g., choanoflagellates, cercozoans, ciliates, centrohelids), putative mixotrophs (e.g., cryptophytes, chrysophytes, dictyochophytes), parasitic protists and fungi. These results demonstrate the potential of qSIP to resolve trophic interactions at fine taxonomic resolution in natural communities and highlight new opportunities to study complex microbial food webs across environmental systems.

## Introduction

In aquatic ecosystems, protists contribute to primary production, form symbiotic associations and connect multiple trophic levels through the microbial loop (Azam et al. 1983; Caron et al. 2017). A major challenge in microbial ecology is linking specific uncultivated taxa, known only via their environmental gene sequences, to their ecological function (Orsi 2023). This extends to heterotrophic protists, whose study is hindered by their large diversity in both lifestyle and evolutionary origin, as well as by prevalent difficulties in cultivation (Campo et al. 2014; Burki et al. 2020). For instance, cultivated representatives and protists that appear abundant in amplicon-sequencing surveys may not necessarily correspond to the most active bacterivores in natural environments (Massana et al. 2009; Rodríguez-Martínez et al. 2022; Wilken et al. 2023). Furthermore, grazing measurements often reflect bulk grazing rates without taxon-specific resolution (Worden et al. 2015), or target only groups for which specific probes are available (Piwosz et al. 2021). As a result, the community-wide identity of protists actively grazing in natural environments remains largely unresolved.

Stable isotope probing (SIP) is a culture-independent technique that links microbial identity and activity to biogeochemical processes and community interactions, by tracing the incorporation of stable isotopes in artificial ratios into the nucleic acids of organisms that have consumed a labeled substrate (Radajewski et al. 2003; Neufeld et al. 2007). SIP has been applied across diverse environments to detect active grazers and infer trophic modes of uncultivated taxa. Early applications identified cercozoan grazers in soil microbial communities using DNA– and RNA-SIP (Lueders et al. 2004) and subsequent RNA-SIP studies revealed ciliated and flagellated grazers of *Escherichia coli* in wastewater and activated sludge (Kuppardt et al. 2010; Moreno et al. 2010). In the open ocean, RNA-SIP has uncovered diverse heterotrophic and mixotrophic protist grazers of the picocyanobacteria *Prochlorococcus* and *Synechococcus* (Frias-Lopez et al. 2009; Wilken et al. 2023), while DNA-SIP has been used to detect grazers of the picoeukaryote *Micromonas pusilla* (Orsi et al. 2018). However, such investigations remain scarce, and to date only two studies have combined SIP grazing experiments with high-throughput sequencing (i.e., Orsi et al. 2018; Wilken et al. 2023), and these have been conducted in marine systems.

Quantitative SIP (qSIP) is an adaptation of SIP that allows quantification of ^13^C or ^15^N assimilation from a labeled substrate into rRNA genes at the operational taxonomic unit (OTU) level (Hungate et al. 2015; Koch et al. 2018). Prior to qSIP, DNA-SIP methods would typically target the densest fractions of DNA gradients, which could be shown to contain an isotopically enriched pool of nucleic acids derived from stable isotope replacement (“Heavy-SIP”; Youngblut et al. 2018). However, since DNA density increases with GC content, these fractions may also include unlabeled, GC-rich DNA. In qSIP, multiple density fractions are sequenced and quantitative PCR (qPCR) of a target gene is used to normalize OTU abundances across the DNA gradients. This approach reduces biases related to taxon GC content, the proportion of isotope-assimilating organisms and enrichment levels, offering greater sensitivity than traditional “Heavy-SIP” approaches (Youngblut et al. 2018). qSIP has been widely applied to studies of prokaryotes (Orsi 2023), but it has not yet been applied to quantify carbon assimilation by uncultivated protists.

Here, we aimed to use qSIP to characterize the diversity of actively feeding protists in two natural freshwater assemblages, a mesotrophic lake and its main inlet stream. Despite a certain degree of homogenizing dispersal, inlet streams often support higher protist richness than the downstream pelagic environment (Crump et al. 2012; Papadopoulou and Lindström 2026). We therefore asked whether this difference in taxonomic richness is also functionally reflected, specifically whether the more diverse inlet community also hosts a broader set of actively grazing protist taxa than the lake. In addition, in such hydrologically connected habitats, dispersal from the inflowing stream is expected to result in a subset of microbial taxa occurring in both locations, so we also aimed to identify both habitat-specific and shared actively feeding protist OTUs in the two locations.

To address this, we applied qSIP combined with 18S rRNA gene qPCR and amplicon sequencing for the first time in a grazing experiment, where live ^13^C, ^15^N-labeled bacterial cells were introduced to natural protist communities. As prey, we used *Limnohabitans planktonicus*, a ubiquitous bacterium found mainly in non-acidic freshwater habitats (Šimek et al. 2010; Jezbera et al. 2013). *Limnohabitans* rapidly responds to algal-derived organic substrates (Pérez and Sommaruga 2006), followed by strong grazing pressure from flagellates (Šimek et al. 2006, 2014), making it a suitable model prey for studying bacterial-protist interactions and carbon flow to higher trophic levels in lake ecosystems (Šimek et al. 2013; Grujčić et al. 2015). Our findings revealed a diverse assemblage of active protist ^13^C, ^15^N-incorporators in both aquatic habitats, including heterotrophic and mixotrophic predators as well as parasitic taxa, highlighting the capacity of qSIP to uncover food web linkages of individual, largely uncultivated, protists.

## Materials and methods

In this experiment, natural microbial communities from an inlet stream and a pelagic site of the same lake were incubated for 36 h with ^13^C, ^15^N-labeled *L. planktonicus* cells as bacterial prey. Incorporation of labeled biomass into protist DNA was used to identify actively grazing taxa using qSIP combined with 18S rRNA gene qPCR and amplicon sequencing.

### Study sites and sample collection

Sampling was conducted at lake Siggeforasjön, a dimictic forest lake approximately 30 km northwest of Uppsala, Sweden. The lake has a surface area of 0.76 km^2^ and a water residence time of 180 days (Brunberg and Blomqvist 1998, p. 738). Water samples were collected from two locations: the main inlet stream (59.9773° N, 17.1279° E) and a pelagic site near the stream outlet (maximum depth: 4.5 m; 59.9771° N, 17.1372° E). Sampling was carried out on 5 September 2024 under low-flow conditions, which were presumed to allow increased protist grazing in the lotic habitat. At the pelagic site, integrated water samples were collected from the upper 2 m of the water column using a Ramberg tube. In the inlet stream, water was collected by submerging a bottle at the water surface, while minimizing sediment disturbance. Due to spatial heterogeneity in depth and in situ dissolved oxygen (DO) concentrations, inlet samples were collected from multiple points along a 5 m stretch in the middle of the stream. A total of 35 L of water was collected from each location and filtered on-site through a 150 *μ*m plankton mesh to remove larger zooplankton.

Upon return to the laboratory, water from each site was transferred into separate 40 L glass aquaria in a temperature-controlled climate room to homogenize the field-collected subsamples. The room was maintained at 18 °C, based on the mean in situ water temperature of the sampling locations, and a photoperiod of 15 h light / 9 h dark was established to simulate natural conditions. Light intensity was set to 3000 lux, corresponding to an approximate photon flux of 160 *μ*mol m⁻² s⁻¹, as measured with a QSL-100 spherical sensor (Biospherical Instruments). Water was stored in the aquaria for one day prior to the start of the experiment to allow organisms to acclimate to the new abiotic conditions and photoperiod, and to provide protist predators time to feed on the natural bacterial communities.

### Physicochemical parameters

In situ measurements of water temperature, DO and its saturation were taken at both sampling locations using a portable probe (HQ40d multi-parameter meter, HACH). A vertical profile of these parameters was also recorded at the pelagic site to assess the stratification status, and the same measurements were repeated after transferring the water to the aquaria. Haziness, as an estimate of turbidity, was assessed in triplicate for both sample types with water collected from the aquaria: Approximately 20 mL of each sample was filtered through a GF/F filter (0.7 μm, diameter 25 mm; Whatman) and the filtrate was collected in clean glass vials. Absorbance was measured at 436 nm using a 5 cm cuvette in a GENESYS 50 UV-Vis spectrophotometer (Thermo Scientific). Haziness was calculated by subtracting the absorbance of the filtered sample from that of the unfiltered one and dividing by the pathlength (Goedkoop and Sonesten 1995). In addition, model-estimated daily data on total water flow at the lake outlet for 2024 were obtained from the S-HYPE model through the Vattenwebb service of the Swedish Meteorological and Hydrological Institute (subcatchment SUBID: 8110; Siggeforasjön).

Total organic carbon (TOC), total nitrogen (TN), and total phosphorus (TP) concentrations were measured from the aquaria and every experimental bottle (see below). Samples were collected in acid-washed 50 mL Falcon tubes, pre-treated via sequential submersion in three baths: deionized water (DI; 24 h), 1.85% HCl (4 h) and DI (24 h). For TOC and TN, subsamples were transferred to 40 mL glass vials, acid-washed (DI for 24 h; 3.7% HCl for 24 h; DI for 24 h) and combusted in a muffle oven (4 h, 450 °C). These samples were stored at 4 °C in the dark until analysis (within five days). TP samples were stored at −20 °C until analysis (within two months). TOC, TN and TP concentrations were measured following previously described protocols (Papadopoulou et al. 2025).

### Culture media

Two media were prepared for the cultivation of the prey bacterium, *L. planktonicus* strain II-D5 (Gram-negative rods, about 0.3–0.4 *μ*m in diameter and 0.9 *μ*m long; Kasalický et al. 2010). NSY medium, based on Hahn et al. (2003) and formulations from the German Collection of Microorganisms and Cell Cultures (DSMZ), was used to cultivate unlabeled *L. planktonicus* cells. For labeling of bacterial prey, ISOGRO-^13^C,^15^N powder with 99 atom % ^13^C and 98 atom % ^15^N (Merck), consisting of lysates from algae cultured with stable isotopes, was used at a concentration of 4 g L⁻¹. Details on media composition, preparation and cell harvesting are provided in the Supporting Information, Culture media and prey harvesting, and Tables S1–S3. Cultures were inoculated from a frozen stock of *L. planktonicus*, and to ensure complete incorporation of stable isotopes, cells used in the experiment were grown through four consecutive culture batches prior to use. The relative abundance of stable isotopes in the labeled culture (^12^C/^13^C, ^14^N/^15^N) was measured through isotope ratio mass spectrometry at the Tandem laboratory, Uppsala University, Sweden (see Supporting Information, Isotopic composition of *Limnohabitans* cells).

### Stable isotope probing experimental setup

A replicated SIP incubation experiment was conducted using 18 autoclaved (121 °C, 15 min) 2 L Schott Duran bottles, each sealed with Duran venting membrane screw caps (0.22 *μ*m) to allow sterile gas exchange. For the two different sites, three experimental conditions were prepared in biological triplicate:

1. Experimental controls: no prey addition.
2. Unlabeled controls: addition of unlabeled *L. planktonicus* cells.
3. ^13^C, ^15^N-prey experiments: addition of ^13^C, ^15^N-labeled *L. planktonicus* cells.

For each of the three conditions above, three bottles were applied and therefore served as biological replicates for each treatment. Prior to bottle preparation, water from each aquarium was gently mixed using a sterile serological pipette and 2 L of stream or lake water were dispensed into each bottle. The experiment was initiated by adding prey at the midpoint of the dark period (t_0_), at a final density corresponding to approximately 20% of the bacterial abundance in the bottles. The concentration of naturally occurring bacteria in the bottles was determined via flow cytometry (see below). The incubation lasted 36 hours (t_36_) to allow sufficient time for prey consumption and protist growth, enabling stable isotope incorporation into protist DNA during genome replication. This duration also minimized potential cross-feeding effects on re-mineralized ^13^C, ^15^N material (e.g., ^15^NH_3_ or ^13^C-dissolved inorganic carbon) resulting from the labeled prey.

### Flow cytometry

Bacterial abundance in the aquaria and prey cultures was measured by flow cytometry to estimate the number of *L. planktonicus* cells required to reach the target prey density in the experimental bottles. In addition, bacterial and protist abundances as well as the proportion of feeding protists were quantified in all experimental bottles every 12 h post-inoculation. All cytometric measurements were performed with a CytoFLEX flow cytometer (Beckman Coulter) equipped with violet (405 nm) and blue (488 nm) lasers for excitation. Fluorescence emission was detected using 525/40 (FL1) and 690/50 (FL3) filters (blue excitation), and 450/45 (FL5) filter (violet excitation). Side scatter (SSC) was recorded with the blue laser.

Bacterial particles were stained with SYBR Green I nucleic acid stain (1×; Thermo Fisher Scientific) and incubated in the dark at room temperature for 15 min. Bacterial populations were identified based on SSC and green fluorescence (FL1). For protists, particles were stained with both SYBR Green I (1×) and LysoSensor Blue DND-167 (LS; 1 *µ*M final concentration, Invitrogen), and samples were incubated in the dark for 10 min. LS acts as a food vacuole stain and has been used to detect feeding microeukaryotes (Carvalho and Granéli 2006; Florenza et al. 2024). Protist populations were distinguished on the FL3 and FL1 channels. Feeding populations were further gated based on FL1 and LS fluorescence (FL5). A single gate was applied uniformly across all samples to differentiate bacterial or protist cells from background signals and particle concentrations were calculated per volume. A “feeding ratio” was calculated per sample as the ratio of LS-stained protists divided by the total protist abundance in that sample. Additional details are provided in the Supporting Information, Flow cytometry.

### DNA extractions

Cells from the initial microbial communities in the aquaria (t_0_) and from all experimental bottles at the end of the incubation (t_36_) were collected on both 0.22 *μ*m and 3 *μ*m filters (47 mm; Cytiva Whatman, Nuclepore polycarbonate track-etched membranes), targeting bacterial and eukaryotic communities, respectively. Additionally, cells from both labeled and unlabeled *L. planktonicus* cultures were filtered through 0.22 *μ*m membranes to assess the presence of the cultured OTU within natural communities. All filters were stored at −70 °C until processing. DNA was extracted using the DNeasy PowerSoil Pro Kit (Qiagen) following the manufacturer’s protocol.

### Quantitative stable isotope probing

Samples for qSIP were collected at the end of the incubation period (t_36_) from all bottles containing ^13^C, ^15^N-labeled prey (*n* = 3 bottles per treatment, per site) and the controls with unlabeled cells (*n* = 3 bottles per treatment, per site). In addition, DNA was extracted from both labeled (*n* = 1) and unlabeled (*n* = 1) *L. planktonicus* cultures used in the experiment to confirm the complete labeling of prey via density gradient ultracentrifugation. Extracted DNA was prepared for ultracentrifugation and density gradient fractionation following established qSIP protocols (Coskun et al. 2024; Trejos-Espeleta et al. 2024) with minor modifications as detailed below.

Extracted DNA (0.5–6 *μ*g) was mixed with cesium chloride (CsCl, 7.16 M, 0.22 *μ*m filtered) and autoclaved gradient buffer (0.1 M Tris, 0.1 M KCl, 1 mM EDTA) to achieve a final density of 1.660 g mL⁻¹. A total of 3.3 mL of this mixture was loaded in OptiSeal polypropylene centrifuge tubes (Beckman Coulter). Density gradient centrifugation was performed using an Optima MAX-TL ultracentrifuge (Beckman Coulter) with a near-vertical rotor at 65,000 rpm (≈1.5 × 10^5^ × g) for 72 h at 18 °C. Immediately after ultracentrifugation, gradients were fractionated from the bottom of each tube using a syringe pump (flow rate: 14.8 mL h⁻¹) and a fraction recovery system (Beckman), yielding 20 fractions of approximately 165 *μ*L each.

Control gradients were fractionated before the labeled samples to minimize cross-contamination from labeled DNA. The refractive index of each fraction was measured immediately after collection (from least to most dense) using an AR200 digital refractometer (Reichert) and converted to density using a standard curve. For the bacterial culture samples, the same procedure was applied, except that the starting density was 1.700 g mL⁻¹.

DNA from all gradient fractions was precipitated overnight at room temperature by adding two volumes of PEG solution (30% polyethylene glycol 6000 and 1.6 M NaCl; sterile-filtered through a 0.22 *μ*m membrane filter and autoclaved) and 4 *μ*L of linear polyacrylamide (LPA). The use of LPA as a carrier was applied, as it greatly reduces contamination compared to glycogen (Bartram et al. 2009). Samples were centrifuged using an Allegra X-30 centrifuge (Beckman Coulter) equipped with a 3015 rotor at 15 °C and 16,000 rpm (≈ 28,672 × g) for 15 min. Supernatants were discarded and pellets were washed with cold 70 % ethanol, followed by a second centrifugation under the same conditions. Ethanol was removed and the pellets were subsequently air-dried at room temperature and resuspended in 30 *μ*L of DEPC-treated water.

Finally, qPCR was performed on all gradient fractions recovered after ultracentrifugation to generate density curves for each sample and its corresponding control. For the inlet and lake incubations, qPCR targeted the protist 18S rRNA genes using primers V4F/V4RB (Balzano et al. 2015). For the *L. planktonicus* cultures, the bacterial 16S rRNA was quantified via qPCR using primers 515F/806R (Pichler et al. 2018). All reactions were conducted on a CFX Connect Real-Time PCR Detection System (Bio-Rad) using the SsoAdvanced Universal SYBR Green Supermix. Detailed qPCR protocols are provided in Tables S4–S5, along with information about the standard curves (Supporting Information, Quantitative PCR).

Weighted average DNA densities for each gradient were calculated as in Hungate et al. (2015). For the experimental bottles, labeled and control gradients processed together in the same batch for the ultracentrifugation, fractionation and qPCR steps were treated as pairs, resulting in three pairs per site. For the *L. planktonicus* cultures, a single labeled and control gradient were analyzed.

### Preparation of amplicon libraries

DNA extracts intended for bacterial 16S rRNA gene amplicon sequencing were submitted for library preparation and sequencing to the National Genomics Infrastructure (NGI; Stockholm, Sweden). Amplification was performed using the 341F/805R primer pair (Hugerth et al. 2014), with phasing nucleotides inserted into both forward and reverse primers of the first PCR (PCR1). For eukaryotic 18S rRNA gene sequencing, the same primers that were used for qPCR were employed. To enhance cluster detection on the Illumina flow cell during the initial sequencing cycles, four random nucleotides (NNNN) were added to the 5’ end of the forward primer targeting the 18S rRNA gene. Primer sequences, including phased variants, are listed in Table S6.

Total community and qSIP-fractionated samples targeting the 18S rRNA gene were amplified in triplicate PCR reactions (thermocycling conditions in Table S7), with technical replicates pooled following amplification. For each density gradient, 7–15 density fractions were sequenced, which provides sufficient resolution to calculate taxon-specific excess atom fraction (EAF) isotope incorporation (Sieradzki et al. 2020). Amplicons were purified using magnetic beads (MagSi-NGSPREP Plus, magtivio) and DNA concentrations were quantified with PicoGreen assays (Quant-iT PicoGreen dsDNA Assay Kit, Invitrogen). All PCR1 products were normalized to a DNA concentration of 1 ng *μ*L⁻¹ prior to submission to NGI for indexing and sequencing. Only qSIP density fraction samples yielding sufficient DNA were included in library preparation. For both 16S and 18S rRNA gene pools, unique dual indices were used for indexing. Sequencing was performed on the Illumina NextSeq 2000 P1-600 (2×300bp), using one flow cell for the 16S rRNA gene and two for the 18S rRNA gene library.

### Processing of raw reads

Primer sequences were removed from the raw reads using Cutadapt (v4.0 in Python 3.9.5; Martin 2011). Quality filtering was performed using the DADA2 pipeline (v1.28.0; Callahan et al. 2016) in R (v4.3.1). OTU tables were generated using the VSEARCH pipeline (v2.30.0; Rognes et al. 2016). Pre-filtered paired-end reads were merged with a minimum overlap of 16 bp for 16S rRNA and 10 bp for 18S rRNA sequences. Subsequent steps included dereplication, clustering at 97% sequence similarity, removal of singletons (i.e., sequences with an abundance < 2), chimera detection and removal. These steps resulted in one OTU table per gene.

Taxonomic classification was conducted using DADA2 with a minimum bootstrap confidence threshold of 80. The SILVA reference database (v138.2; Quast et al. 2013; Yilmaz et al. 2014) was used as the taxonomy training set for 16S rRNA sequences, while the PR2 reference database (v5.1.0; Guillou et al., 2012) was used for 18S rRNA sequences. The resulting 18S rRNA OTU table was further curated using LULU (lulu package v0.1.0; Frøslev et al. 2017). Contaminant sequences were removed prior to downstream analyses. For 16S rRNA, chloroplast, mitochondrial and unclassified sequences at the phylum level were excluded. For 18S rRNA, sequences assigned to domain “Bacteria”, those unclassified at the domain level and supergroup levels were removed. Sequences from non-protist or non-grazer organisms (e.g., Metazoa, Streptophyta) were intentionally retained to assess the performance of qSIP in distinguishing true grazers from potential false positives. All further analyses were conducted in R (version 4.4.2) and Rstudio (v2024.9.1).

### Alpha and beta diversity

Rarefaction curves were generated using the *rarecurve* function from the vegan package (v2.6-10; Oksanen et al. 2025). For selected analyses, datasets were rarefied without replacement using the *rarefy_even_depth* function from phyloseq (v1.50.0; McMurdie and Holmes 2013). Alpha diversity was estimated as observed richness (i.e., number of OTUs) from the rarefied datasets. For the 16S rRNA, differences in richness between stream and lake whole (non-fractionated) communities were assessed using a Wilcoxon rank sum test (*wilcox.test*; stats package, v4.4.2) due to the non-normal distribution of the data. For the 18S rRNA, a Welch two-sample t-test (*t.test*; stats) was applied. To visualize changes in microbial community composition, OTU counts from non-rarefied datasets were transformed into relative abundances. These were summarized at the phylum level for 16S rRNA and subdivision level for 18S rRNA, for both whole community and fractionated samples.

Community composition based on rarefied 16S and 18S rRNA gene datasets was analyzed using non-metric multidimensional scaling (NMDS) based on Bray-Curtis dissimilarity matrices. NMDS was performed using the *ordinate* function (phyloseq) with Bonferroni correction. To assess the homogeneity of group dispersions (i.e., the distance of samples from their group centroid), the *betadisper* function (vegan) was used. Differences in dispersion between groups were then tested using a permutation-based test (*permutest*; vegan) with 999 permutations. To evaluate whether microbial communities clustered significantly by sampling site, permutational multivariate analysis of variance (PERMANOVA) was performed using the *adonis2* function (vegan), with 9,999 permutations and Bonferroni correction.

### Quantitative taxon-specific ^13^C and ^15^N incorporation

Protist taxon-specific changes in DNA density due to isotope incorporation were calculated following Hungate et al. (2015) and applying two additional criteria (hereafter “qSIP criteria”):

1. OTUs had to be detected in all three bottle replicates, of both the labeled and control incubations, at each site.
2. OTUs were required to have at least 50 reads in a minimum of five gradient fractions per bottle incubation. This threshold was chosen to ensure the formation of complete OTU-level density curves as using fewer fractions (e.g., three) often resulted in incomplete normal distributions across the density gradient, where only one side of the expected bell-shaped distribution was formed.

Additional modifications based on Hungate et al. (2015) were applied as follows:

1. For each OTU, the buoyant density shift (*Z*) at t_36_ was defined as the difference in weighted average DNA density between an isotopically labeled incubation and its corresponding paired control (i.e., the control processed in the same ultracentrifugation and fractionation batch).
2. Since both carbon and nitrogen isotopes were simultaneously labeled in the prey culture, we estimated a combined enrichment value by weighing the isotope specific EAF values according to their relative contributions to DNA mass, assuming carbon contributes approximately 2.5 times more than nitrogen (Angel 2019). The EAF values accounted for the background fractional abundance of ^15^N as 0.003663004 (Finley et al. 2019). The EAF represents the quantitative amount of isotope incorporation into protist 18S rRNA genes and ranges from 0 to 1, where 1 corresponds to complete (100%) replacement of carbon and nitrogen in the DNA with ^13^C and ^15^N derived from the labeled prey cells. We then calculated the bootstrap mean EAF (100 resamples) across the three replicate pairs of labeled and control incubations, along with 99% confidence intervals (CIs). OTUs were considered potential incorporators of labeled prey if their lower CI did not overlap zero.

We subsequently assessed the presence of unique and shared actively feeding OTUs across sites. Rank abundance curves were also generated for the initial inlet and lake rarefied 18S rRNA gene communities at t_0_ by assigning each OTU a unique rank. Labeled OTUs from each site were plotted on these curves to assess whether labeled taxa were restricted to abundant ranks or also included rare taxa.

### Taxonomic refinement of qSIP OTUs

To refine the taxonomic annotations initially obtained through DADA2 classification against the PR2 database, all OTUs meeting the qSIP criteria (both labeled and unlabeled) were queried against the NCBI Core Nucleotide Database using BLASTn. A top-ranking hit was selected for each OTU based on sequence similarity and annotation quality. Taxonomy was assigned as either (1) the lowest PR2 rank (excluding species rank) with a bootstrap support ≥ 95 in DADA2, or (2) the genus rank for BLAST alignments with species annotation at sequence identity > 99 % and E-value < 1e^-150^.

SSU rRNA gene phylogenies were also performed to refine functional annotations (pigmentation, feeding behavior) when assigned taxonomy was unclear. First, a selection of PR2 reference sequences (v5.1.2) was clustered to 99 % similarity (except Eustigmatophyceae, clustered to identity) using VSEARCH (v2.22.1) to reduce redundancy and excessive taxon sampling. The Pseudofungi phylogeny was supplemented with additional sequences from Prokina et al. (2024). The resulting sequence sets (File S1) were subsequently used to build alignments with the L-INS-i algorithm in MAFFT (v7.526; Katoh and Standley 2013), to which query OTUs were added a posteriori. Final alignments were trimmed with trimAl (v1.5.rev1; Capella-Gutiérrez et al. 2009), imposing a 30 % gap threshold. Maximum likelihood phylogenies were constructed with IQ-TREE (v3.0.1; Wong et al. 2025) using the General Time Reversible (GTR) nucleotide substitution model. Taxonomic annotations were thus amended when phylogenies resolved deeper taxonomic detail at > 95 bootstrap support.

## Results

### Environmental conditions

Model-estimated daily total water flow at the outlet of lake Siggeforasjön indicated high discharge during winter and spring and low flow during summer and early autumn, with sampling conducted during the driest period of 2024 (Fig. S1). A weak temperature gradient (<1 °C m⁻¹) was observed throughout the lake water column, indicating the breakdown of thermal stratification prior to autumnal mixing (Table S8). Following water transfer to the aquaria, DO concentrations increased slightly for both sites, reaching 7.43 mg L⁻¹ (74 % oxygen saturation) for the inlet stream and 9.56 mg L⁻¹ (97.4 %) for the lake water. TOC, TN and TP concentrations were consistently higher in the stream than in the lake incubations (Table S9). On average, TOC, TN and TP concentrations in the inlet were 24 mg L⁻¹, 1.0 mg L⁻¹ and 170 *μ*g L⁻¹ (*n* = 10), respectively, compared to 15 mg L⁻¹, 0.43 mg L⁻¹ and 13 *μ*g L⁻¹ in the lake (*n* = 10). The inlet was also 49 times hazier (0.343 ± 0.003 cm^-1^) than the lake water (0.007 ± 0.0002 cm^-1^; Fig. S2).

### ^13^C, ^15^N-labeling of *Limnohabitans* cells

When grown in isotopically labeled medium, the isotopic composition of *L. planktonicus* cells, as determined by isotope ratio mass spectrometry, was 66.7 ± 3.0 atm% (mean ± standard deviation [SD]) for ^13^C and 77.9 ± 2.2 atm% for ^15^N (*n* = 11). These values likely underestimated the true isotopic enrichment, as the ^13^C, ^15^N-labeled cells had to be diluted with unlabeled material prior to analysis due to their high degree of labeling (*cf.* qSIP results for *L. planktonicus* cultures).

### Bacterial and protist abundances

Prior to setting up the SIP incubations, bacterial abundance was 2.4 × 10^6^ cells mL⁻¹ in the inlet sample and 3.8 × 10^6^ cells mL⁻¹ in the lake sample. Based on these values, 9.4 × 10^8^ prey cells (corresponding to the 20 % target) were added to the inlet incubations and 1.5 × 10^9^ cells to the lake incubations at t_0_, both contained in 2 L bottles. After 12 h post-prey addition, bacterial abundance in the inlet water exceeded that in the lake, and at both sites a decreasing trend in cell densities was observed from 12 h until the end of the experiment at 36 h under all three experimental conditions (Fig. 1a). Protist abundances were consistently higher in the inlet than in the lake, but showed no clear temporal trend or consistent differences among experimental conditions (Fig. 1b). In contrast, the feeding ratio, calculated as the LS-stained particles divided by the total protist abundance, was consistently lower in the inlet than in the lake and remained relatively stable over time (Fig. 1c). We note that the high haziness of the inlet samples may have affected the overall accuracy of flow cytometric measurements. Consequently, the number of prey cells added to the stream incubations at t_0_ may have been less precisely quantified compared to the lake samples, and may have deviated from the intended 20 % target. Nevertheless, the prey concentration added was sufficient to induce detectable shifts in protist density, as revealed by qSIP (see below).

**Figure 1.**
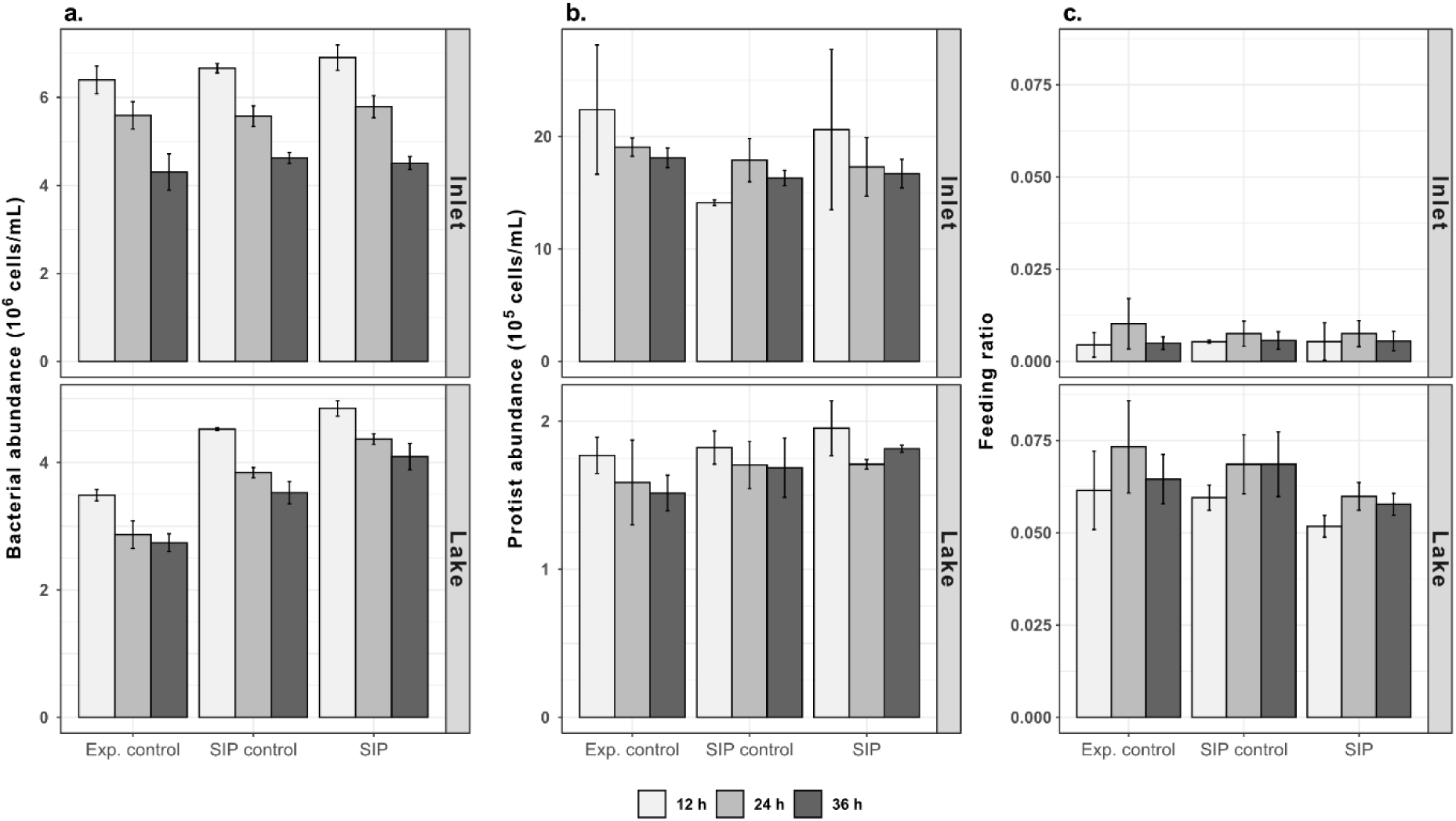
Flow cytometry measurements in inlet stream and lake experimental bottles from 12 h post-prey addition to 36 h (end of the experiment). (a) Bacterial abundance, (b) protist abundance, (c) feeding ratio. Colors indicate the time points measured. Samples are grouped by experimental condition: experimental controls without prey addition (“Exp. control”), unlabeled controls (“SIP control”) and ^13^C, ^15^N-prey experiments (“SIP”). Bars represent mean values across replicate incubations (*n* = 3); error bars indicate ± standard deviation. Note that panels a and b use different y-axis scales for inlet and lake samples.

### Community composition and diversity

Community structure differed significantly between inlet and lake environments for both bacterial and eukaryotic communities, as shown by NMDS ordinations (stress < 0.05 for both; Fig. S3) and PERMANOVAs (16S rRNA whole communities: F_1,18_ = 145.13, *p* < 0.001; 18S rRNA whole and fractionated communities: F_1,139_ = 154.87, *p* < 0.001; Table S10). The 16S rRNA dataset contained 11,079 OTUs. After rarefaction to 508,072 reads per sample, 10,705 OTUs were retained. The 18S rRNA dataset contained in total 2,735 OTUs; qSIP-fractionated communities contained 2,660 OTUs. Following rarefaction to 377,024 reads per sample, 2,560 OTUs remained. For both bacteria and eukaryotes, whole communities from inlet samples exhibited significantly higher richness than those from the lake (16S rRNA: Wilcoxon rank-sum test, W = 100, *p* < 0.001; 18S rRNA: Welch two-sample t-test, t = 13.80, df = 11.44, *p* < 0.001; Fig. S4). Rarefaction curves for whole and fractionated communities are provided in Figures S5 and S6.

At the beginning of the experiment (t_0_), inlet and lake communities shared 14.95 % of the total bacterial OTUs and 30.94 % of the total eukaryotic OTUs (Fig. 2). For both bacteria and eukaryotes, the inlet community contained the highest number of OTUs, followed by the shared fraction, while the lake community contained the fewest OTUs. Bacterial communities at t_0_ were dominated by the phyla Pseudomonadota, Actinomycetota and Bacteroidota in both the lake and the inlet. A higher relative abundance of Patescibacteria was observed in the inlet, whereas higher relative abundance of Planctomycetota and Verrucomicrobiota was observed in the lake. As per PR2 classification, eukaryotic communities were dominated by the subdivision Gyrista in both environments. The inlet was enriched in sequences from Cercozoa, Choanoflagellata and Ciliophora, while the lake showed higher relative abundances of Cryptophyta, Dinoflagellata and Metazoa. After 36 h of incubation, the relative abundances of the major bacterial phyla and eukaryotic subdivisions remained relatively stable across treatments in both inlet and lake experimental bottles (Files S2–S3), indicating that the experimental conditions (e.g., “bottle effects”) did not markedly affect community structure. In the freshwater communities and ^13^C, ^15^N-prey cultures, a dominant bacterial OTU (bacterial OTU 1) was identified, affiliated with *Limnohabitans* (> 99 % of reads in the prey cultures, 100 % bootstrap support) which was rare in the inlet community at t_0_ (0.27 % relative abundance), but an abundant member of the initial lake community (26.48 %).

**Figure 2.**
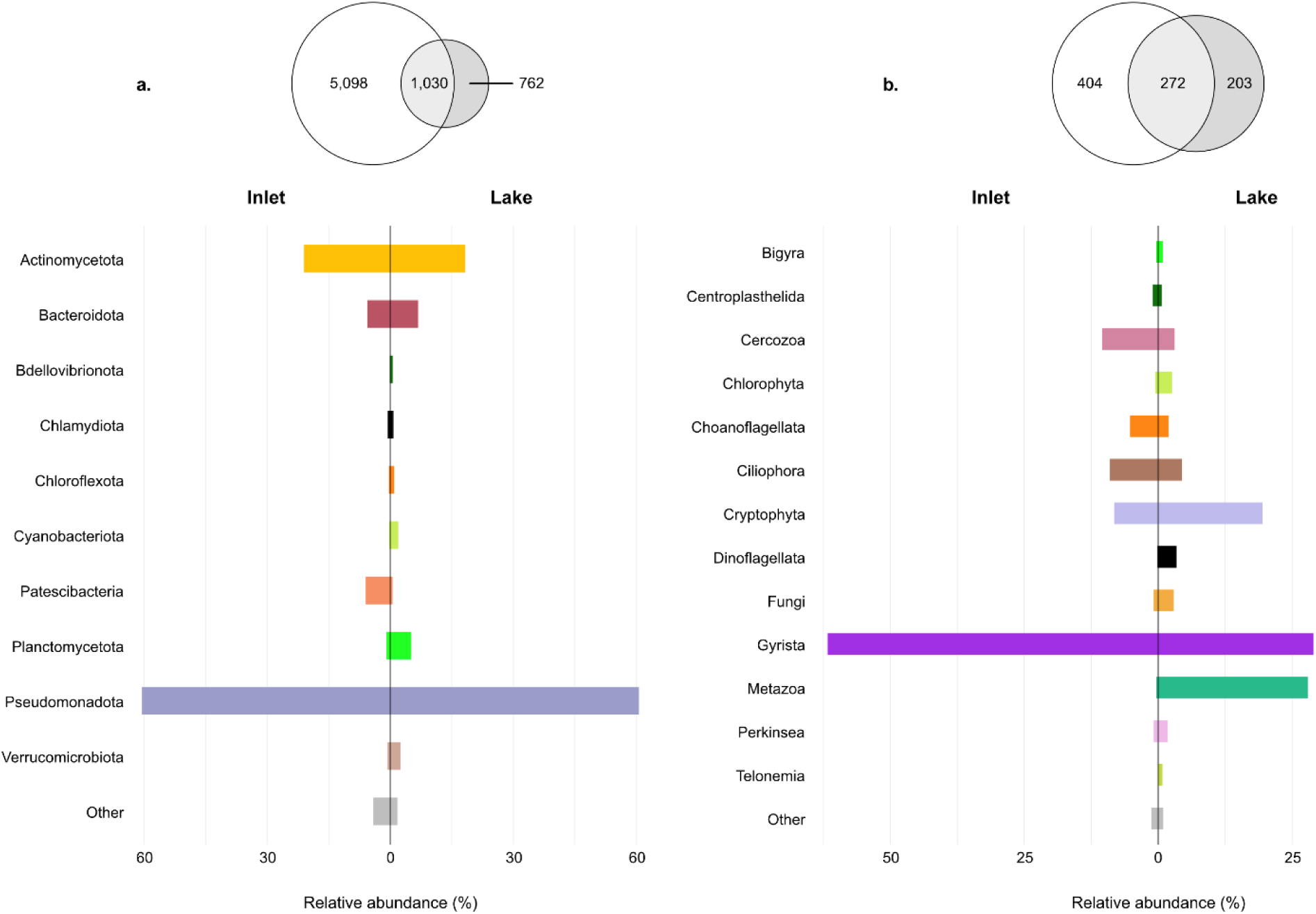
Alpha diversity and taxonomic composition of microbial communities in the initial inlet and lake samples (t_0_). (a) Rarefied bacterial communities assessed via 16S rRNA gene sequencing and (b) rarefied eukaryotic communities characterized using 18S rRNA gene sequencing. The upper panels show Euler diagrams illustrating the number of unique and shared operational taxonomic units (OTUs) between the stream (white circle) and lake (grey circle) environments. The lower panels present bar charts of the relative abundances of dominant bacterial phyla and eukaryotic subdivisions, based on DADA2-derived taxonomy.

### Community-level 18S rRNA gene density shifts

The qPCR-based profiles of 18S rRNA genes showed buoyant density shifts in all biological replicates, indicating ^13^C, ^15^N-isotopic enrichment of protist DNA derived from the labeled prey cells (Fig. 3a, b). In the inlet samples, the mean increase in buoyant density (*Z)* between the labeled and control treatments was 0.0018 ± 0.0020 g mL⁻¹ (mean ± SD; *n* = 3 bottle replicates), while in the lake, a slightly higher *Z* of 0.0058 ± 0.0036 g mL⁻¹ (*n* = 3 bottle replicates) was observed. Figure 3 also shows the fractions that contained sufficient DNA for successful 18S rRNA gene amplification. The relative abundance of eukaryotic subdivisions for all sequenced fractions can be found in File S4. The DNA density gradient centrifugation analysis of *L. planktonicus* cultures showed a complete separation in buoyant density between unlabeled and labeled DNA, indicating near-complete replacement of the ^13^C and ^15^N isotopes into the prey cell DNA (Fig. 3c). Therefore, the modest increase in protist DNA buoyant density can be attributed to the consumption of nearly fully-labeled *L. planktonicus* prey cells that were added to the SIP incubations.

**Figure 3.**
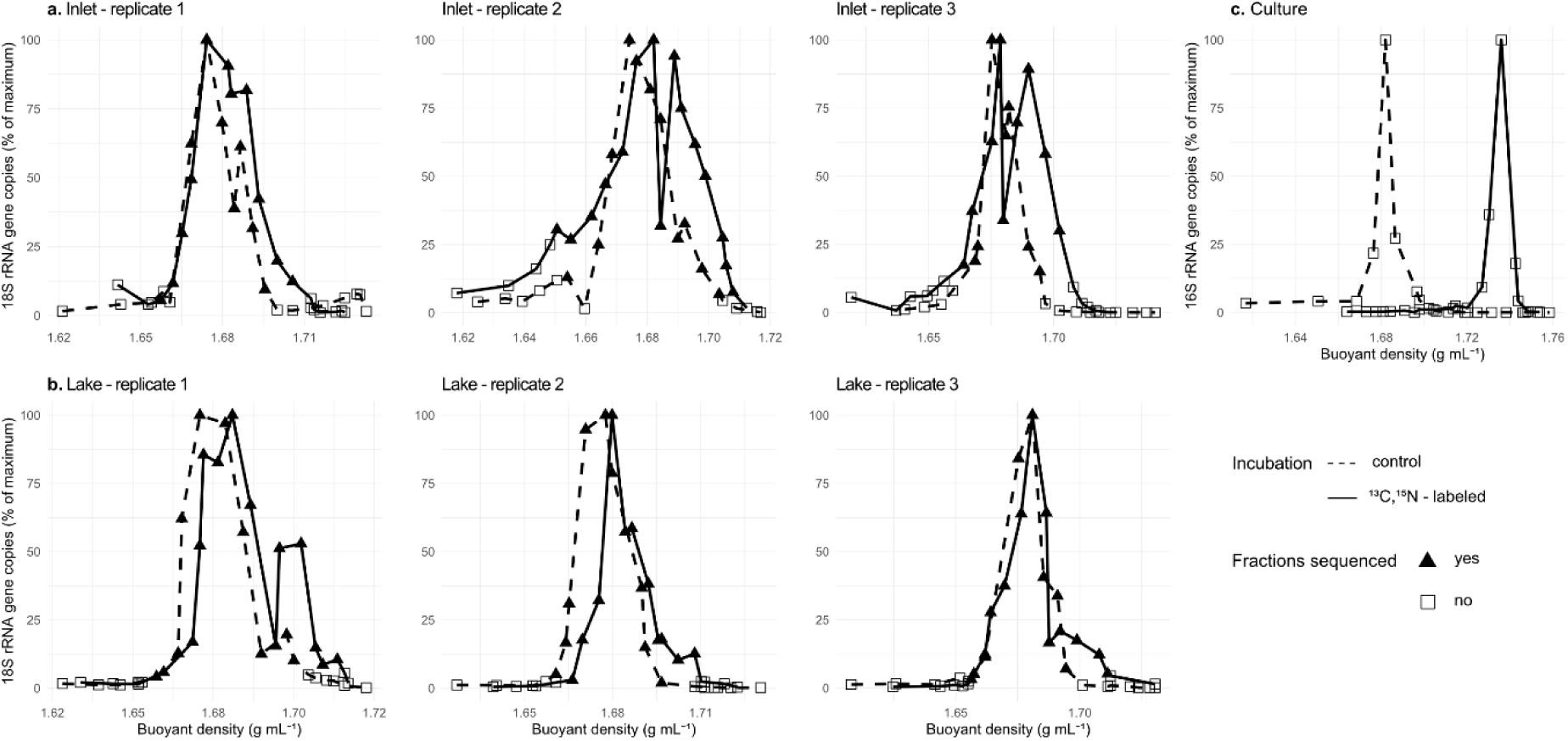
Density gradient profiles of 18S rRNA gene abundance across CsCl density gradients from SIP incubations of the. (a) inlet and (b) lake. Increases in buoyant density in the labeled treatments relative to the unlabeled controls indicate incorporation of ^13^C and ^15^N isotopes into DNA. (c) Buoyant density shift of *L. planktonicus* DNA, demonstrating strong isotopic labeling of the prey cells prior to their use in the grazing experiment. DNA fractions successfully sequenced for 18S rRNA genes are indicated by filled triangles.

### Active isotope incorporators identified by qSIP

Regardless of sample origin, OTU richness in both labeled and unlabeled incubations declined with increasing buoyant density, likely reflecting GC content-driven DNA migration in CsCl gradients (Fig. S7). Inlet samples exhibited higher OTU richness than lake samples in the lighter fractions; however, both sample types showed similar levels of richness in the heavier fractions.

A total of 126 OTUs met the qSIP criteria in the inlet stream (i.e., were detected in all labeled and unlabeled replicate incubations per site and had ≥ 50 reads in at least five gradient fractions) compared with 197 OTUs in the lake, with 49 OTUs shared between sites. Of the total OTUs meeting the qSIP criteria, 108 inlet OTUs and 107 lake OTUs were labeled (i.e., exhibited a positive *Z*, with the lower 99% confidence interval not overlapping zero; Fig. 4–5). Notably, 26 OTUs were active ^13^C, ^15^N-incorporators in both environments. The protist OTU showing the greatest incorporation in the inlet was affiliated with Crustomastigaceae (OTU 235, subdivision Chlorophyta; *Z* = 0.32 g mL⁻¹, 99 % CI: 0.25–0.36), a potential phago-mixotrophic prasinophyte (Fig. 4). In the lake community, the protist with the greatest isotope incorporation was affiliated with Chrysophyceae (OTU 166, subdivision Gyrista; *Z* = 0.65 g mL⁻¹, 99% CI: 0.54–0.73; Fig. 4–5). Labeled protist OTUs from both environments spanned 16 subdivisions according to PR2 taxonomy and included both pigmented and non-pigmented taxa, representing putative heterotrophic, mixotrophic and parasitic trophic modes (Tables 1–2). Many of the labeled OTUs were affiliated with uncultivated lineages.

**Figure 4.**
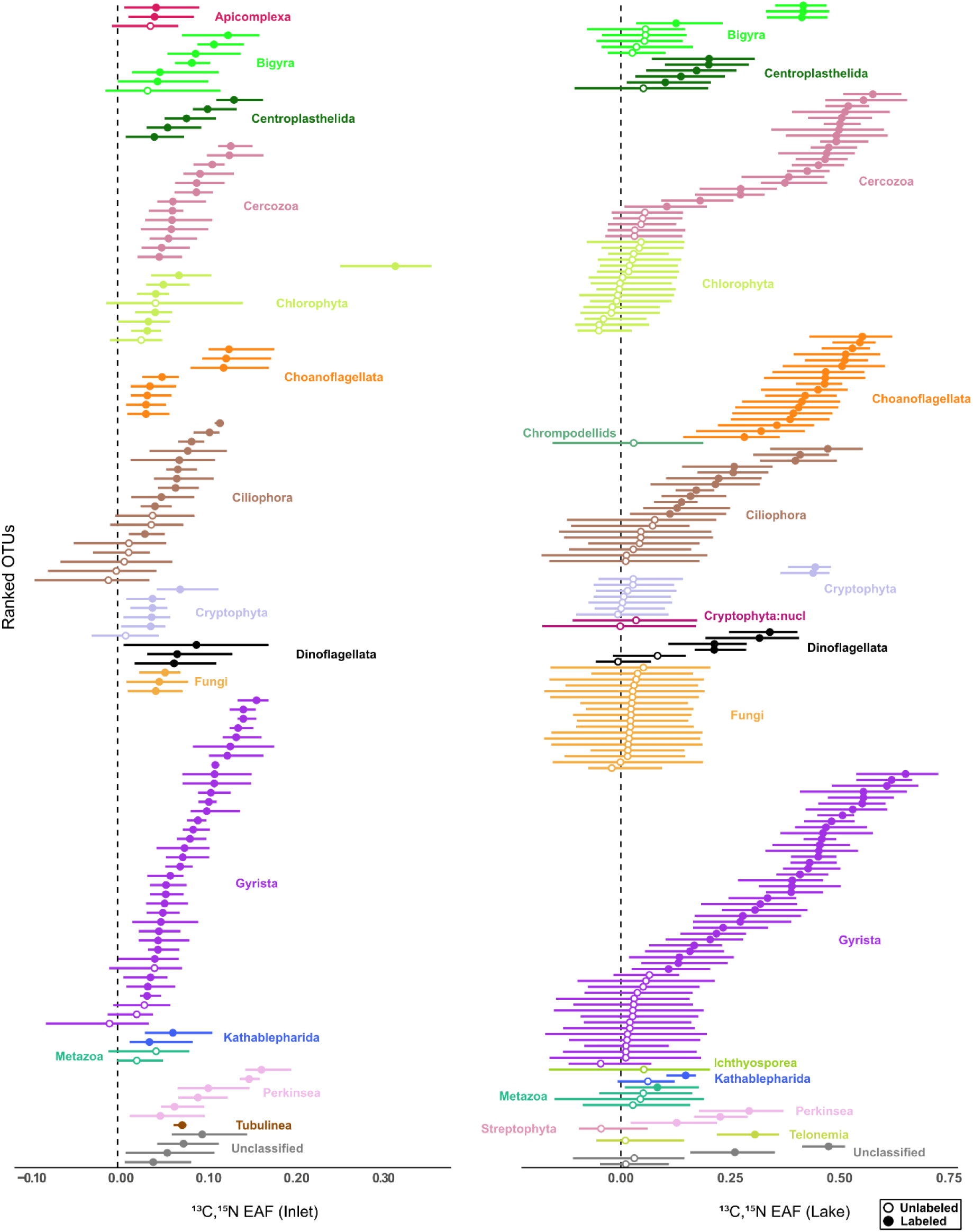
Excess atom fraction (EAF) values for eukaryotic OTUs from the inlet stream (left) and lake pelagic site (right) incubations. OTUs are ranked by mean EAF and grouped by subdivision (as per PR2 taxonomy). Solid circles indicate significant incorporators of labeled *L. planktonicus* prey, while open circles represent unlabeled OTUs whose buoyant density shift did not exceed the significance threshold for isotope incorporation. Protist OTUs were considered significant incorporators of labeled prey when the lower 99% confidence interval (CI) did not overlap zero (99 % CI shown as error bars).

**Figure 5.**
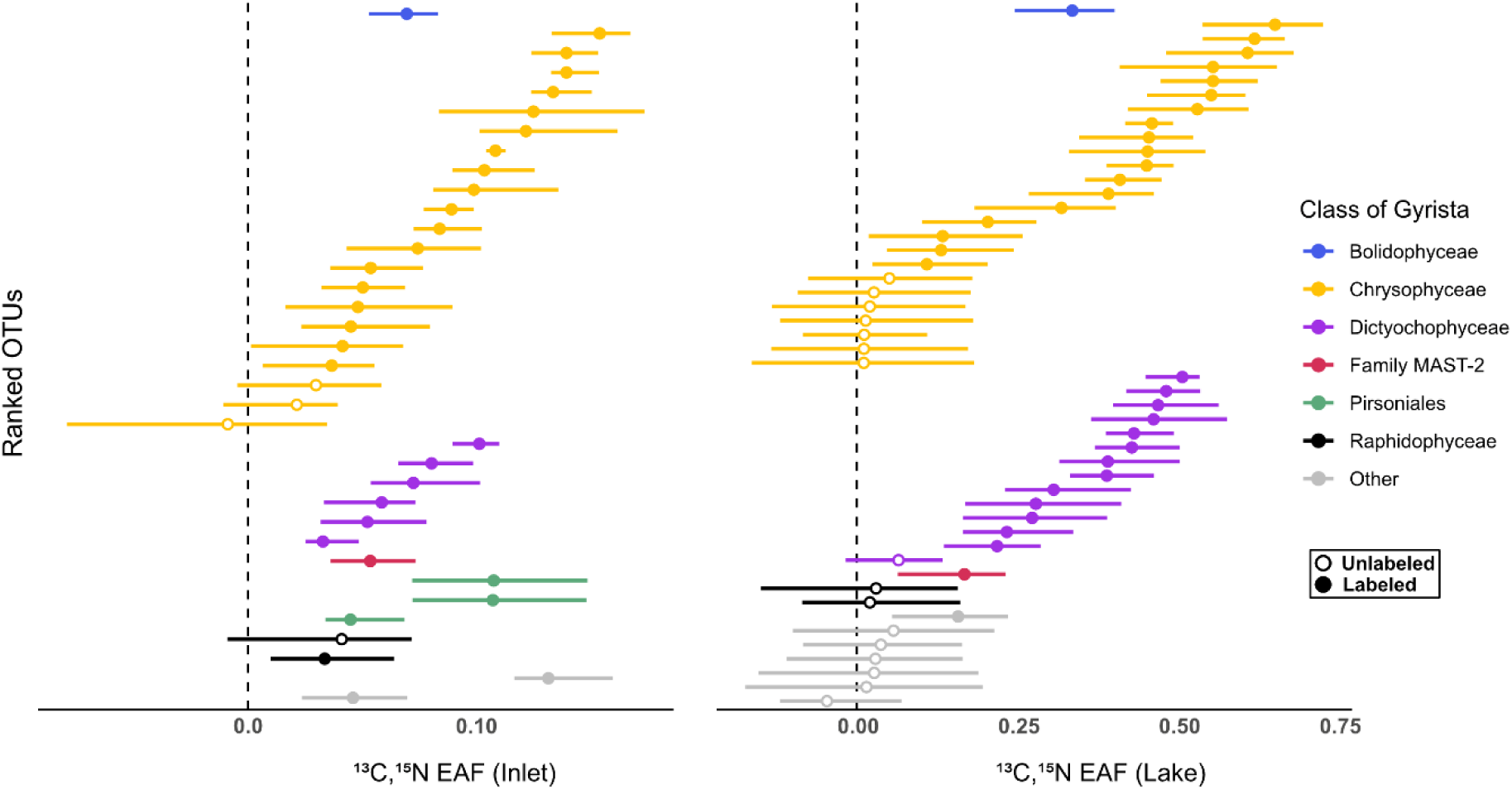
Excess atom fraction (EAF) values for OTUs within the subdivision Gyrista (as per PR2 taxonomy). OTUs are colored by class. This figure presents a more detailed view of the Gyrista data shown in Figure 4. Solid circles indicate active incorporators of labeled prey.

**Table 1.**
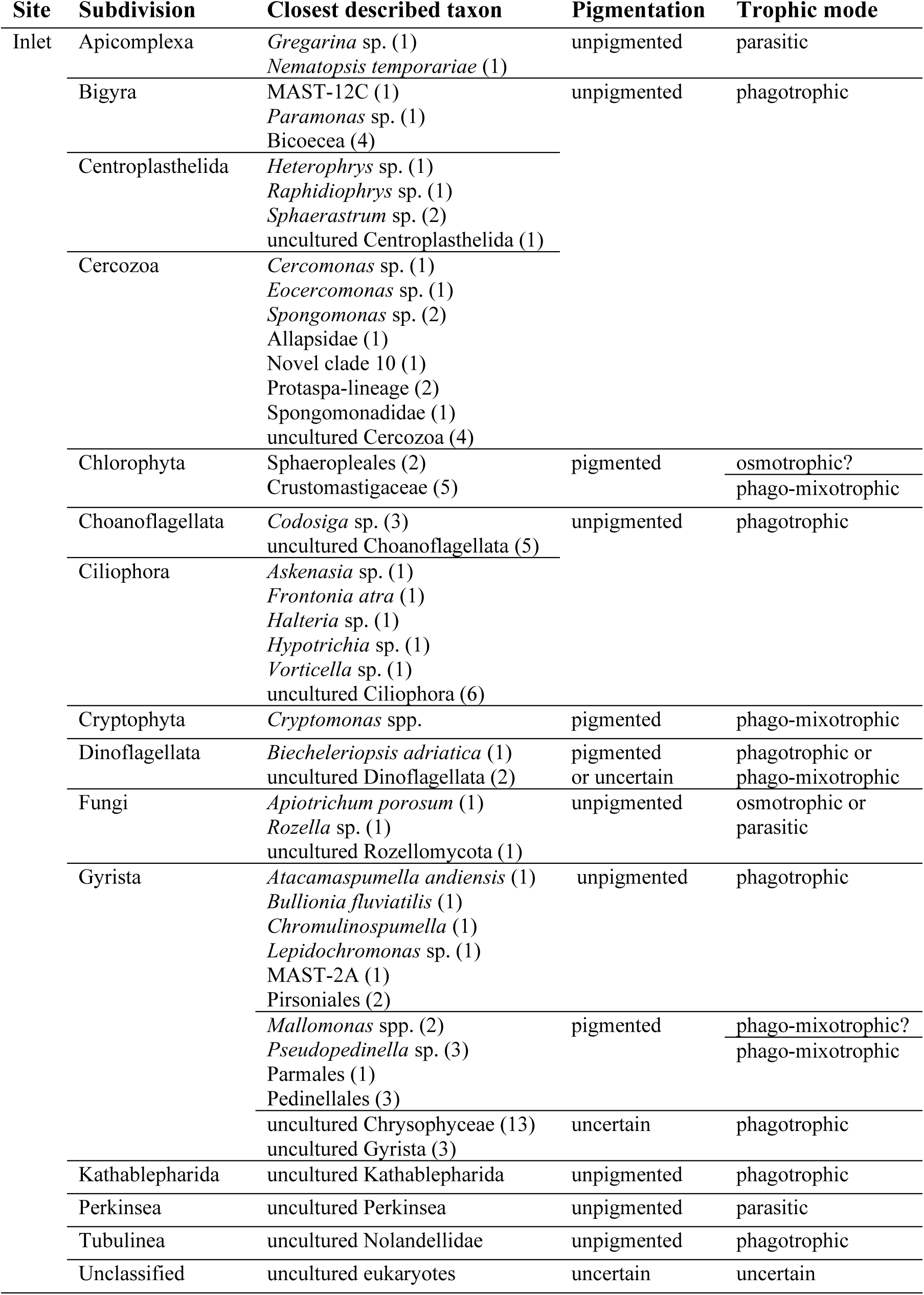
Taxonomic summary of operational taxonomic units (OTUs; *n* = 108) actively incorporating labeled bacterial biomass in the inlet stream qSIP incubations. The table reports, for each taxonomic group (subdivision, as per PR2): the number of OTUs, the closest described taxon based on curated annotations (see File S5; number of OTUs in parentheses), and the putative pigmentation and trophic mode observed in the experiment.

**Table 2.**
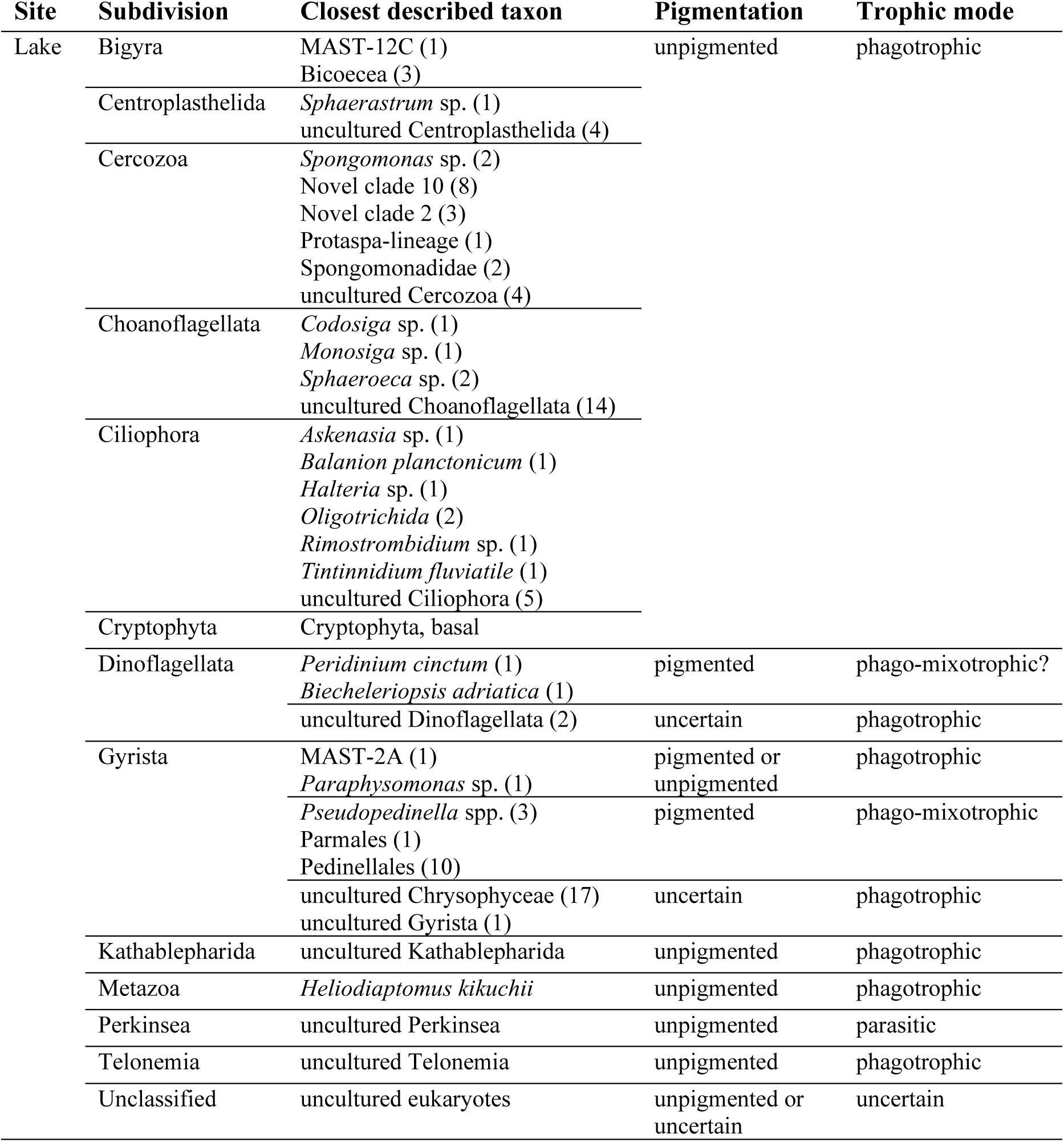
Taxonomic summary of OTUs (*n* = 107) actively incorporating labeled bacterial biomass in the lake qSIP incubations. The table reports, for each taxonomic group (subdivision, as per PR2): the number of OTUs, the closest described taxon based on curated annotations (see File S5; number of OTUs in parentheses), and the putative pigmentation and trophic mode observed in the experiment.

Minimal cross-feeding was suggested by the presence of several unlabeled OTUs that were not expected to feed directly on bacteria. For example, in the lake incubations, all photosynthetic Chlorophyta (*n* = 16), Streptophyta (OTU 308) and a fish parasite from Ichthyosporea (OTU 103) showed no buoyant density shift. Similarly, all OTUs affiliated with Metazoa (e.g., rotifers and copepods) were unlabeled in both inlet and lake incubations, except one lake copepod (OTU 2, *Heliodiaptomus kikuchii*; Z = 0.08 g mL⁻¹, 99% CI: 0.01–0.18). In addition, no fungal OTUs were labeled in the lake incubations (*n* = 18), consistent with their non-phagotrophic, osmotrophic lifestyle. In contrast, some potentially osmotrophic OTUs in the inlet showed relatively low levels of labeling (*Z* < 0.05 g mL⁻¹), including Chlorophyta (*Stauridium tetras*, OTU 200; Sphaeropleales, OTU 120), Gyrista (*Mallomonas* spp., OTU 49 and OTU 99) and Fungi (*Apiotrichum porosum*, OTU 455), likely reflecting minor uptake of dissolved organic matter during the 36-h incubations (Přibyl and Cepák 2019; Peltomaa and Taipale 2020).

File S5 provides a comprehensive compilation of all OTUs meeting the qSIP criteria (labeled and unlabeled), including curated taxonomic, pigmentation and trophic annotations, EAF values (bootstrap mean ± 99 % CI) and information on whether OTUs were shared between inlet and lake incubations. Phylogenetic trees used to refine taxonomic annotations are also provided in the Supporting Information.

Rank abundance curves for the inlet and lake communities at t_0_ showed that ^13^C, ^15^N-labeled OTUs were distributed across a wide range of ranks, indicating that both abundant and rare taxa were recovered as active incorporators (Fig. S8). The least abundant labeled OTUs in both communities had a relative abundance of 0.0024 % at t_0_: OTU 315 (subdivision Bigyra; *Z* = 0.05 g mL⁻¹, 99 % CI: 0.02–0.12) in the inlet and OTU 203 (subdivision Cercozoa; *Z* = 0.47 g mL⁻¹, 99 % CI: 0.36–0.53) in the lake.

## Discussion

In this study, we applied qSIP for the first time in a grazing experiment to investigate the diversity of actively feeding protists in two natural freshwater communities. Using ^13^C, ^15^N-labeled *Limnohabitans* as bacterial prey cells in 36-h incubations of a mesotrophic lake and its main inlet stream, we detected active grazers as organisms showing a positive shift in DNA buoyant density after incorporating prey-derived ^13^C and ^15^N into their DNA. This approach allowed us to link prey assimilation directly to OTU-level taxonomic identity and revealed a diverse assemblage of more than 100 labeled OTUs per site, encompassing both heterotrophic and mixotrophic taxa, while also indicating the presence of parasitic interactions within the food webs.

Both bacterial and protist communities differed significantly in structure between the inlet and lake sites, both in situ and in our incubations. Despite being hydrologically connected, the communities shared only 31 % of their eukaryotic taxa at the start of the experiment (t_0_). The inlet also exhibited higher protist richness than the lake, consistent with previous studies, including earlier work in this lake (Crump et al. 2012; Papadopoulou and Lindström 2026). This pattern likely reflects the greater influence of surrounding soils as a source of microbial dispersal into the stream, a process that may be particularly pronounced under dry conditions.

The cultured bacterium used as prey, affiliated with *Limnohabitans* (bacterial OTU 1), was detected in both natural communities. At t_0_, bacterial OTU 1 was rare in the inlet (0.27 % relative abundance), but was a dominant member of the lake community (26 %). Its high relative abundance in the lake aligns with previous observations (Šimek et al. 2010, 2014; Grujčić et al. 2015) and confirms that this model organism was an ecologically relevant choice for studying protist predator-prey interactions in our system. Moreover, given the dominance of *Limnohabitans* in the initial lake community, the protist predators detected here by qSIP likely represent a major link for carbon and nitrogen transfer to higher trophic levels also in situ.

### qSIP detected both abundant and rare protist incorporators

Although the inlet exhibited higher protist richness than the lake, a similar number of OTUs became labeled after 36 h of incubation (inlet: 108 OTUs, lake: 107 OTUs). Of these, 26 OTUs (24 %) actively assimilating prey, were shared between the two sites. However, many taxa remained unlabeled, including groups not expected to feed on bacterial prey, such as photosynthetic protists and fungi. Importantly, qSIP detected labeled OTUs across a wide rank – abundance spectrum, including rare taxa down to 0.0024 % relative abundance at both sites. Previous in silico work has suggested that isotope labeling is mainly detectable in moderately to highly abundant OTUs (> 0.1 % relative abundance; Youngblut et al. 2018). This broad detection was likely enabled by the deep sequencing depth employed in our study. Identifying such rare feeders would be considerably more challenging using microscopy-based approaches, such as catalyzed reporter deposition-fluorescence in situ hybridization (CARD-FISH; Piwosz et al. 2021), highlighting an advantage of qSIP in capturing a wide range of active grazers at the community level.

### Phagotrophic predators of *Limnohabitans*

We recovered a diverse assemblage of bacterivorous protists, both heterotrophic and mixotrophic, among active incorporators of *Limnohabitans* prey. Typical bacterivores in our dataset included bicosoecids, choanoflagellates, cryptophytes and ochrophytes, the last two represented by either pigmented or colorless forms. Cercozoans and ciliates were also prominent among labeled taxa. Although cercozoans are generally considered bacterivorous, several taxa are omnivorous, capable of ingesting both bacteria and protists (Piwosz and Pernthaler 2011; Šimek et al. 2020; Mills et al. 2025). Likewise, many ciliates are omnivorous, including the here-detected *Rimostrombidium* and *Halteria.* Among potentially omnivorous occurrences, highly labeled OTUs likely represented active bacterivores, while lower EAF values may reflect reduced bacterial consumption or indirect labeling via predation on other fast-feeding bacterivores that had consumed *Limnohabitans*, resulting in isotopic dilution. Similarly, labeled katablepharids at both sites and the flagellate *Telonemia* in the lake likely reflect a combination of bacterivory and indirect labeling via predation on labeled protists (Šimek et al. 2020; Boukheloua et al. 2024).

Interestingly, the recovery of several Crustomastigaceae (Chlorophyta) and one bolidophyte sequences as prey incorporators exemplifies the discovery potential of qSIP to uncover unexpected bacterivorous taxa. Bacterivorous Crustomastigaceae are represented by five OTUs found exclusively in the inlet, including the OTU showing the highest buoyant density shift. Recent experimental and model-based evidence from marine prasinophytes indicates that they are capable of bacterivory (Bock et al. 2021; Xiao et al. 2025), although it has been shown that prasinophytes did not feed when heat-killed bacteria or magnetic beads were provided, suggesting a strong preference for live prey (Bock et al. 2021). This highlights the advantage of our grazing experiment using live bacterial cells. Regarding bolidophytes, the same OTU (order Parmales) was labeled in both sites. While bolidophytes are well-known mixotrophs in marine environments (Li et al. 2022; Ban et al. 2023), they are rarely reported from freshwater. This observation represents, to our knowledge, the first evidence of an actively feeding bolidophyte in freshwater habitats.

### **I**sotopic signal detected in parasites

A number of positively labeled OTUs belonged to putative parasitic groups, namely apicomplexan gregarines, perkinsids and fungi. For these taxa, isotopic enrichment can hardly be attributed to direct engulfment of labeled bacterial prey, but rather positive labeling would be the result of parasitic interactions with other eukaryotes that originally fed on labeled prey, either directly or indirectly. For example, it is possible that perkinsids and rozellids (the latter found exclusively in the inlet) acquired enriched carbon and nitrogen indirectly from parasitizing other protists, such as dinoflagellates (Gleason et al. 2012; Reñé et al. 2021; Cooney et al. 2024). Similarly, the recovery of apicomplexan OTUs could reflect parasitism in the gut of crustaceans (Sano et al. 2016). Although parasitic interactions were not the focus of this study, their detection highlights the complex network of trophic interactions among bacteria, protists and parasites that qSIP can reveal, involving both the microbial loop and the mycoloop (Kagami et al. 2014).

### Experimental considerations

Previous studies using *Limnohabitans* as prey focused on simplified microbial food webs by excluding larger protists (> 5–10 *μ*m; e.g., Grujčić et al. 2018; Šimek et al. 2018, 2020). In contrast, we retained larger grazers to capture the full diversity of actively feeding taxa in the inlet and lake communities. Our design did not allow us to distinguish strict bacterivores from omnivores, which tend to be larger (8–15 μm; Šimek et al. 2020; Piwosz et al. 2021). Thus, future experiments could use different size fractions of the protist community or conduct separate incubations with isotopically labeled bacteria and protists to disentangle omnivory from bacterivory.

Despite a similar number of active prey incorporators at both sites, EAF values were generally lower in the inlet incubations (3–32%) than in the lake (8–65%). This likely reflects challenges for protists in locating non-motile *L. planktonicus* cells in the particle-rich inlet water. Achieving buoyant DNA density shifts comparable to those observed in the lake may have required the addition of more labeled cells or longer incubation times. However, our goal was to assess the diversity of actively feeding protists at each site rather than to directly compare incorporation levels, while maintaining labeled substrate concentrations within environmentally relevant ranges. The flow cytometry-based feeding ratio further suggested lower overall grazing activity in the stream. Although comparisons of protist grazing activity between lentic and lotic habitats are scarce, lower ingestion rates in streams have been reported previously (Peters 1994).

Given the high diversity of labeled taxa recovered in both stream and lake environments, grazer taxa appeared to exhibit low prey selectivity. This is further reinforced by the fact that qSIP-detected grazers likely represent only a subset of protist predators capable of consuming *Limnohabitans* across different seasons and locations (Grujčić et al. 2015). The diversity of grazers may in fact have been higher than detected due to primer bias against certain taxonomic groups (e.g., Excavata). Nevertheless, predator preference for the *L. planktonicus* strain used here appeared high, either highlighting the central role of *Limnohabitans* as an ecologically important genus in freshwater habitats or indicating that factors other than prey taxonomy influence grazing (Florenza and Bertilsson 2023). Future qSIP experiments using more complex labeled prey communities could help resolve patterns of prey selectivity.

Overall, this study represents the first application of qSIP in a grazing experiment demonstrating that qSIP can simultaneously, effectively and quantitatively detect a multitude of active bacterivores. We recovered a diverse assemblage of actively feeding protists, including heterotrophic and putative mixotrophic taxa, in two natural freshwater habitats without prior knowledge of community composition. Protist richness was higher in the stream compared to the lake, yet the richness of active grazers detected by qSIP was similar at both sites. The application of qSIP in comparative studies of protist grazing dynamics across freshwater, marine and soil ecosystems could help better link specific protist taxa to bacterial grazing within the highly diverse protist communities found in natural environments.

## Author contributions

**Sofia Papadopoulou:** Conceptualization, Data Curation, Formal Analysis, Funding Acquisition, Investigation, Methodology, Project Administration, Software, Validation, Visualization, Writing – Original Draft Preparation. **Javier Florenza:** Conceptualization, Data Curation, Formal Analysis, Investigation, Methodology, Software, Validation, Visualization, Writing – Review & Editing. **Christoffer Bergvall:** Formal Analysis, Methodology, Validation, Writing – Review & Editing. **Eva S. Lindström:** Conceptualization, Funding Acquisition, Resources, Supervision, Writing – Review & Editing. **William D. Orsi:** Conceptualization, Formal Analysis, Methodology, Validation, Resources, Supervision, Writing – Review & Editing.

## Data availability statement

Raw sequence data have been deposited in the European Nucleotide Archive under accession number PRJEB104077 for the 16S rRNA and PRJEB104079 for the 18S rRNA genes sequences. OTU and taxonomy tables for amplicon sequencing, and flow cytometry data (.FCS files) are available in Zenodo (https://doi.org/10.5281/zenodo.19137998). All other relevant data supporting the findings of this study are available within the article and its supporting information. Custom scripts developed for data analysis are available from the corresponding author upon reasonable request.

## Supporting information

Phylogenetic trees used to refine taxonomic annotations

The resulting sequence sets (File S1)

sequenced fractions can be found in File S4

File S5 provides a comprehensive compilation of all OTUs meeting the qSIP criteria

provided in the Supporting Information, Culture media and prey harvesting

## Acknowledgements

This work was supported by the Swedish Research Council [grant number 2020-03110 to EL], a Research and Training Grant from the Federation of European Microbiological Societies [to SP], and the Deutsche Forschungsgemeinschaft [DFG grant number OR 417/7-1 to WDO]. The authors acknowledge support from the National Genomics Infrastructure in Stockholm funded by Science for Life Laboratory, the Knut and Alice Wallenberg Foundation and the Swedish Research Council, and the National Academic Infrastructure for Supercomputing in Sweden (NAISS) for assistance with massively parallel sequencing and access to the UPPMAX computational infrastructure. The authors also thank Marina Ivanković for providing the *L. planktonicus* strain and Yvonne Meyer-Lucht for assistance in acquiring research materials.

## Conflicts of Interest

None declared.

## Notes

### Competing Interest Statement

The authors have declared no competing interest.

