## Supplementary figures and images for "Protist quantitative stable isotope probing identifies diverse active grazers in natural freshwater communities"

### IQ.Bigyra-v2.gt30.GTR.18S.contree.revised.pdf

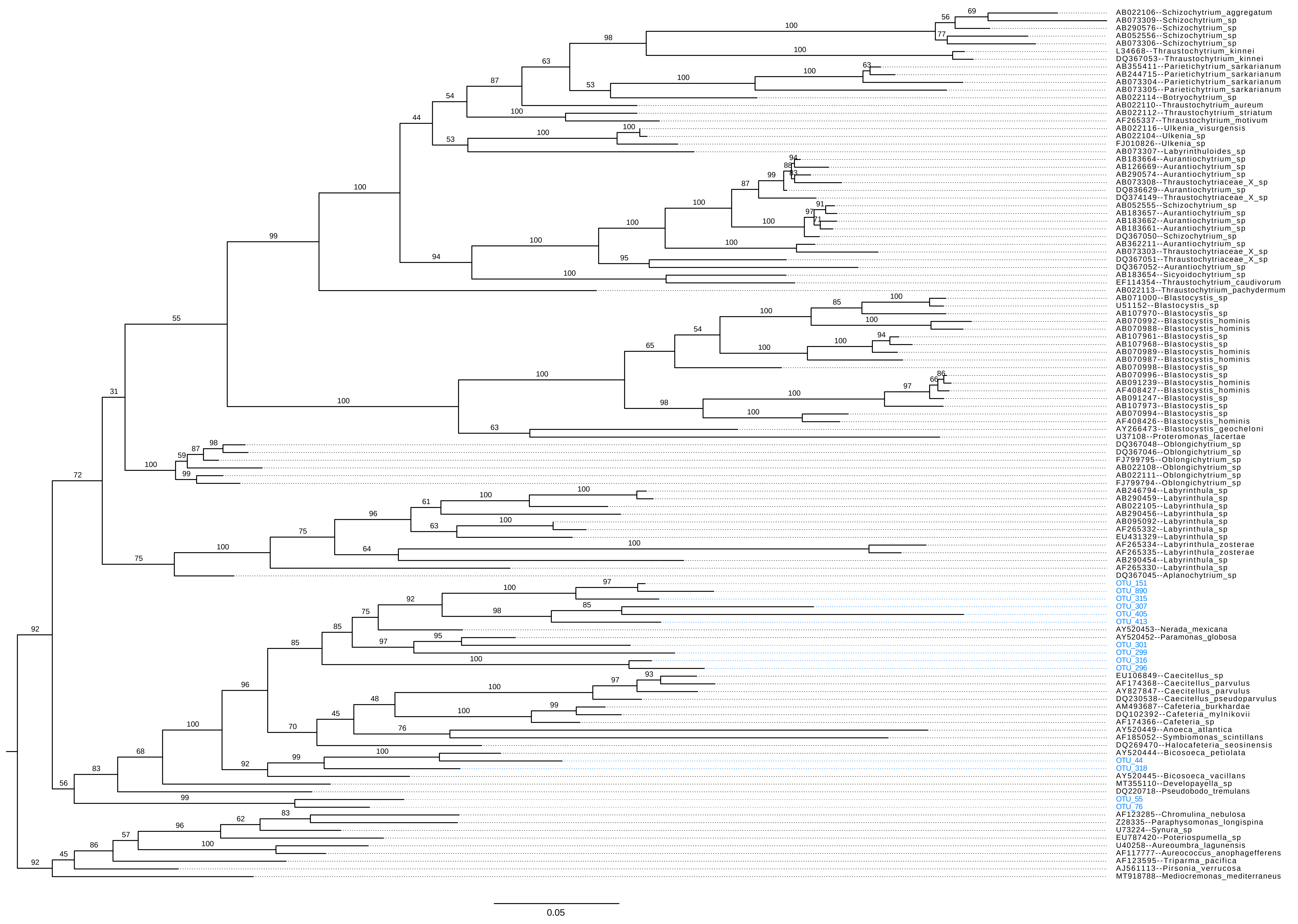

### IQ.Cercozoa-v3.gt30.GTR.18S.contree.revised.pdf

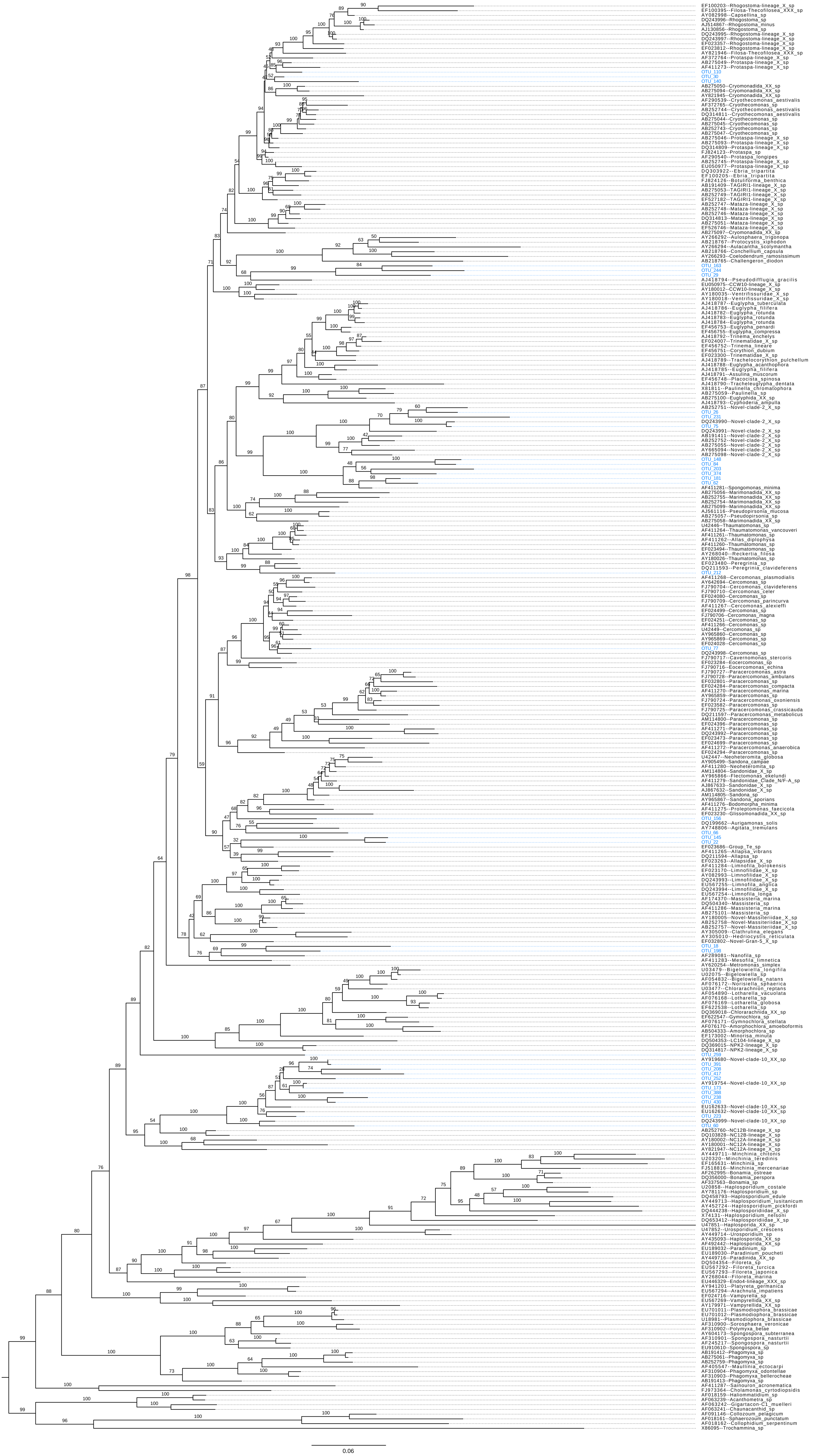

### IQ.Chrysista-v4.gt30.GTR.18S.contree.revised.pdf

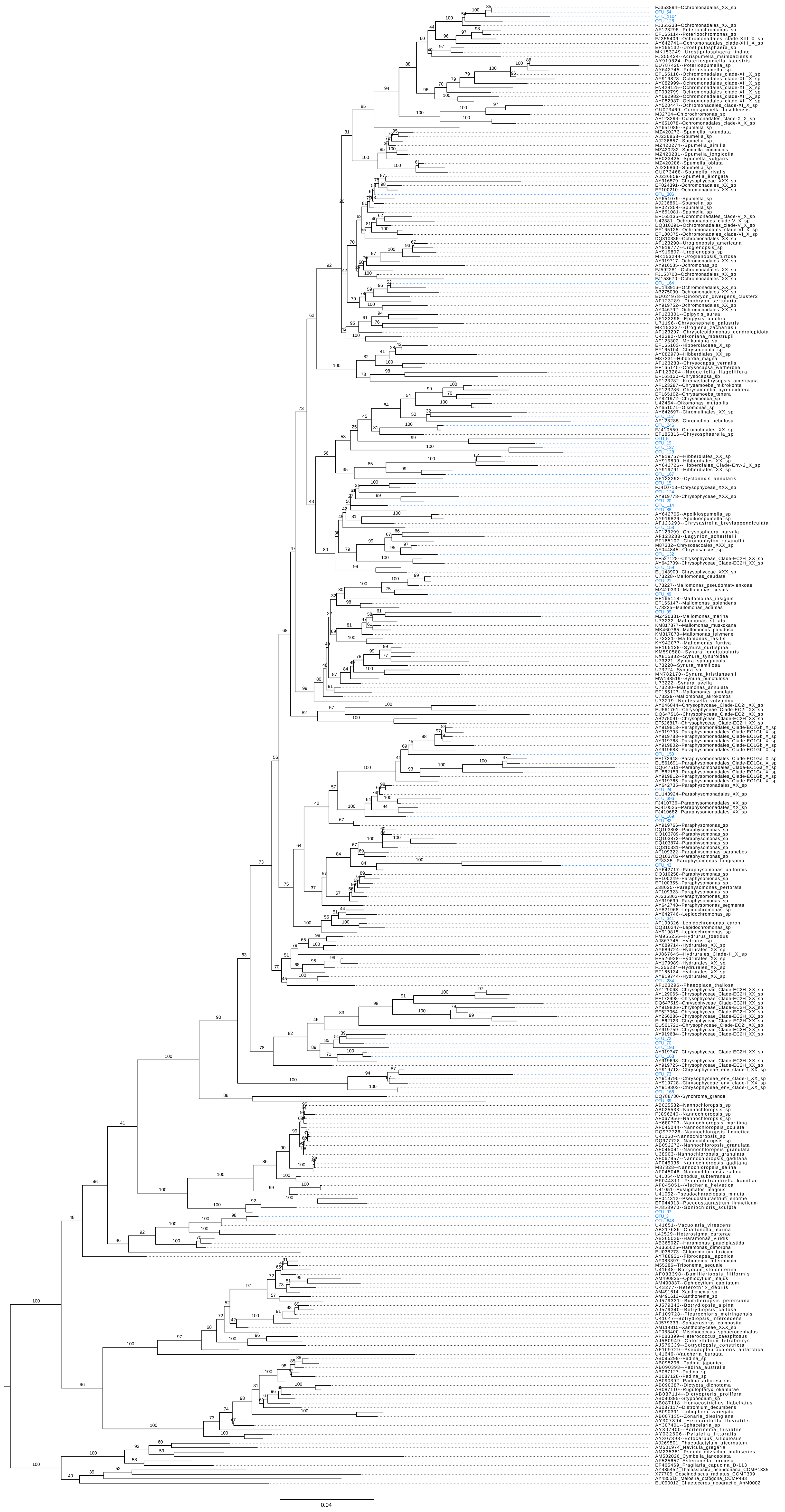

### IQ.Cryptista-v3.gt30.GTR.18S.contree.revised.pdf

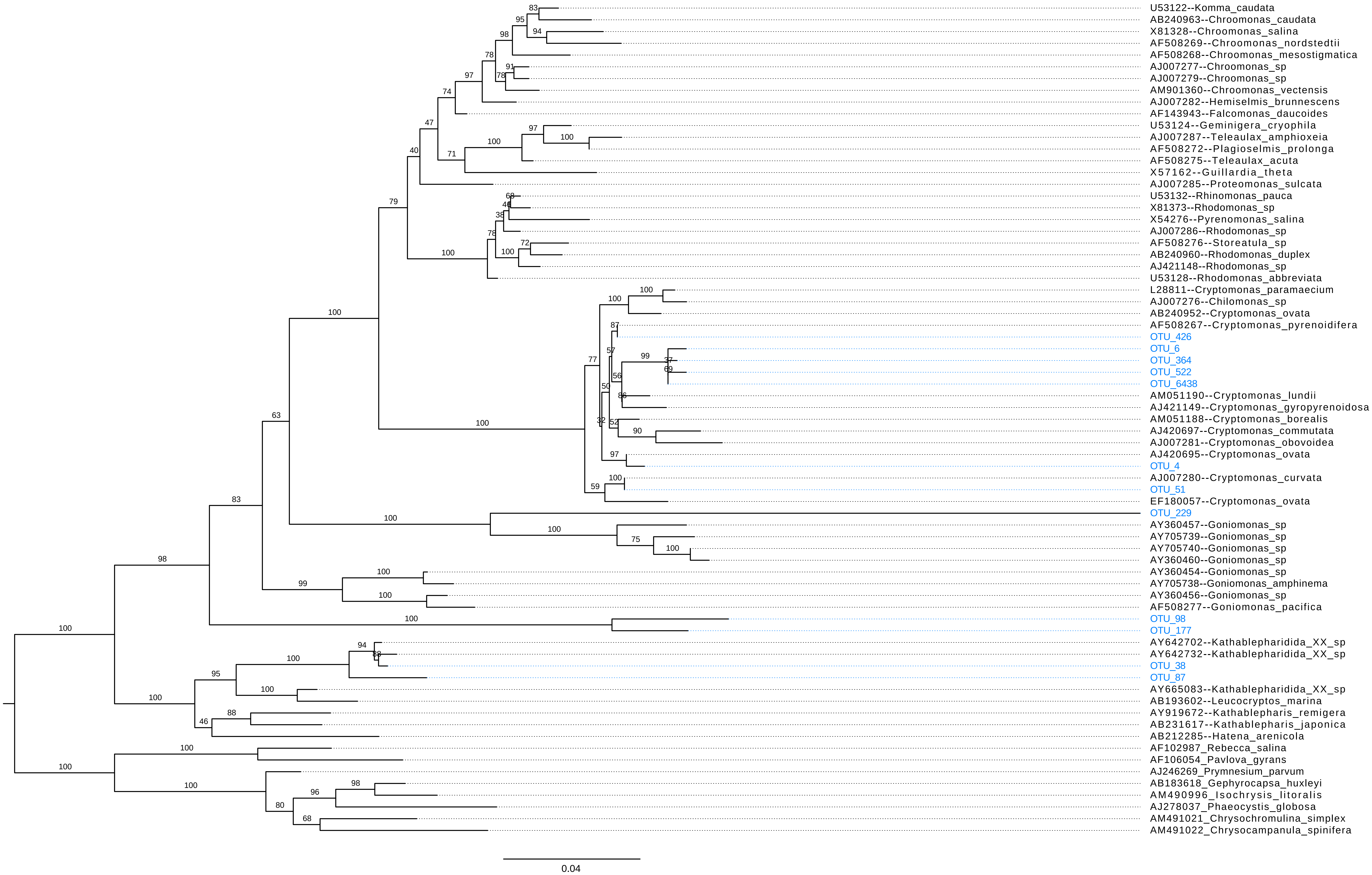

### IQ.Cryptophyta-Nucleomorph.gappyout.GTR.18S.contree.revised.pdf

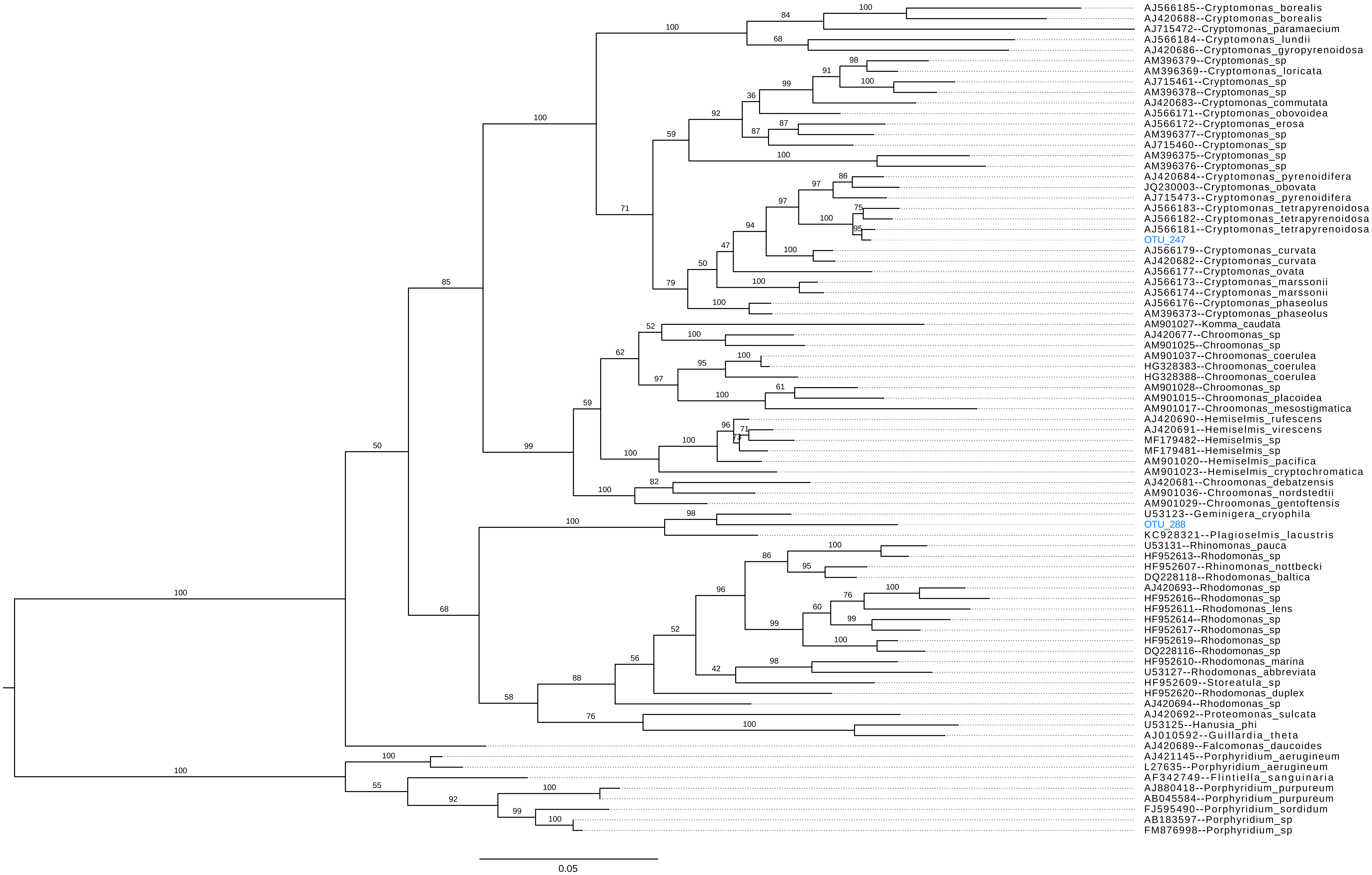

### IQ.Diatomista-v2.gt30.GTR.18S.contree.revised.pdf

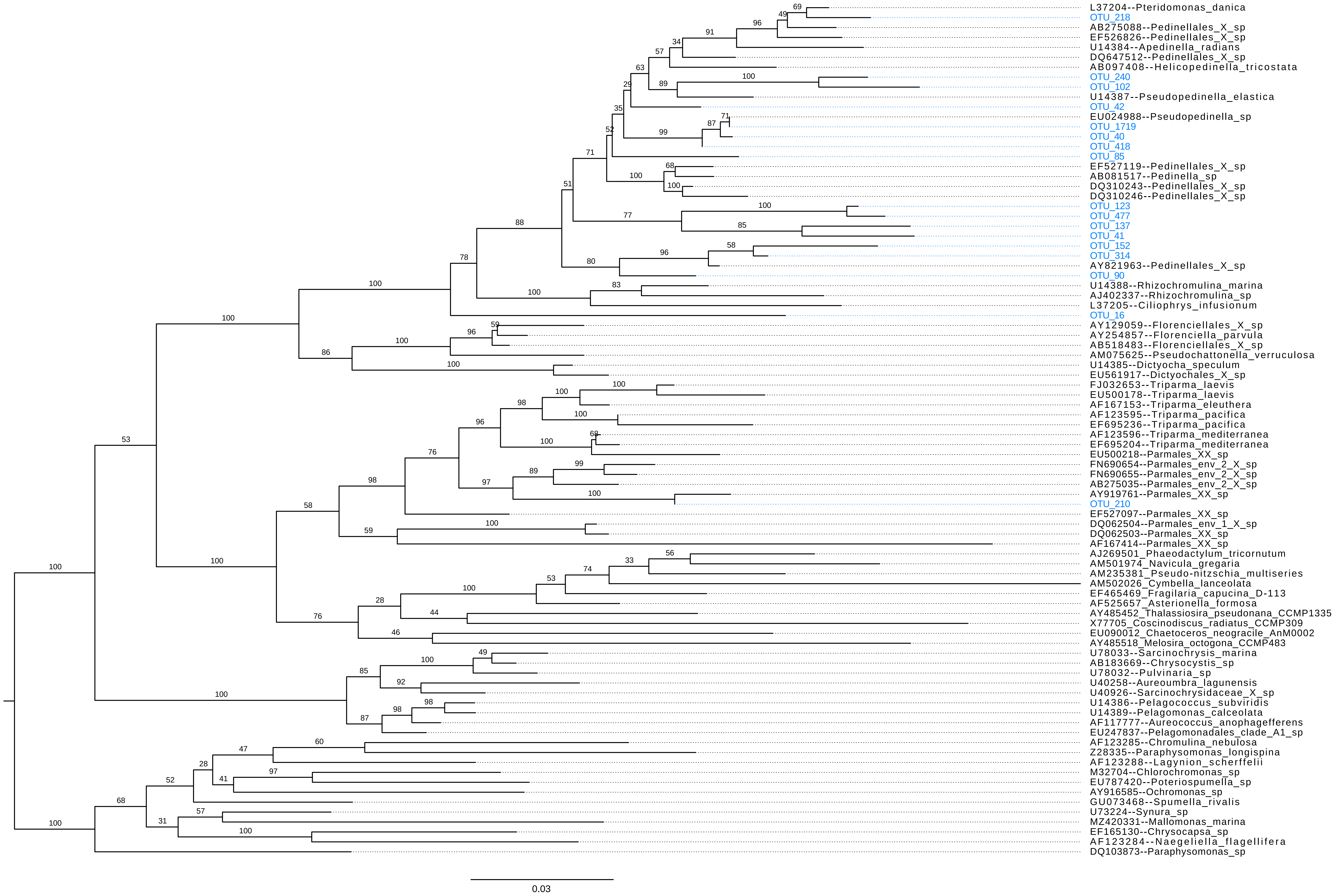

### IQ.Pseudofungi-v2.gt30.GTR.18S.contree.revised.pdf

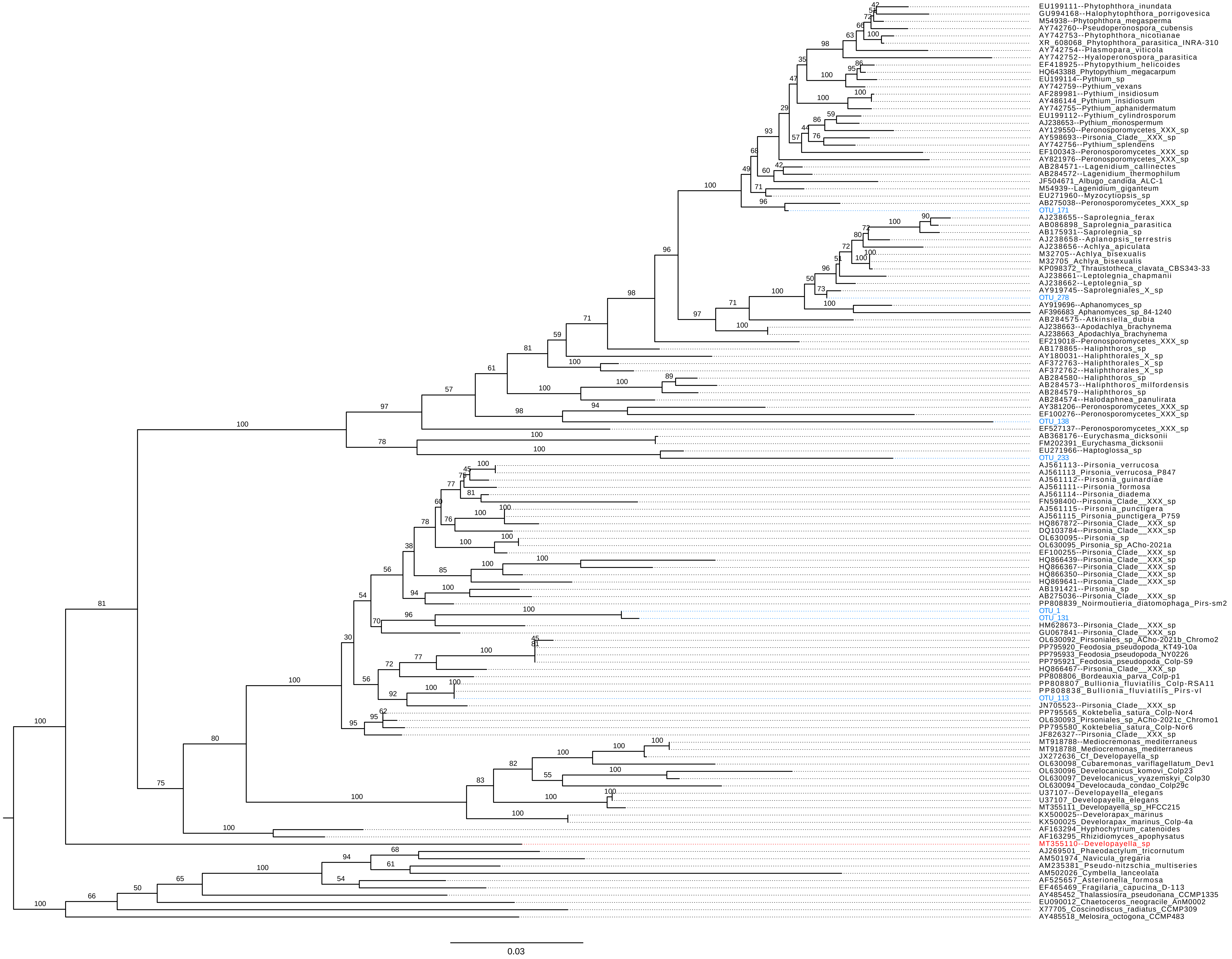
