## Supplemental Data 1 for "Protist quantitative stable isotope probing identifies diverse active grazers in natural freshwater communities"

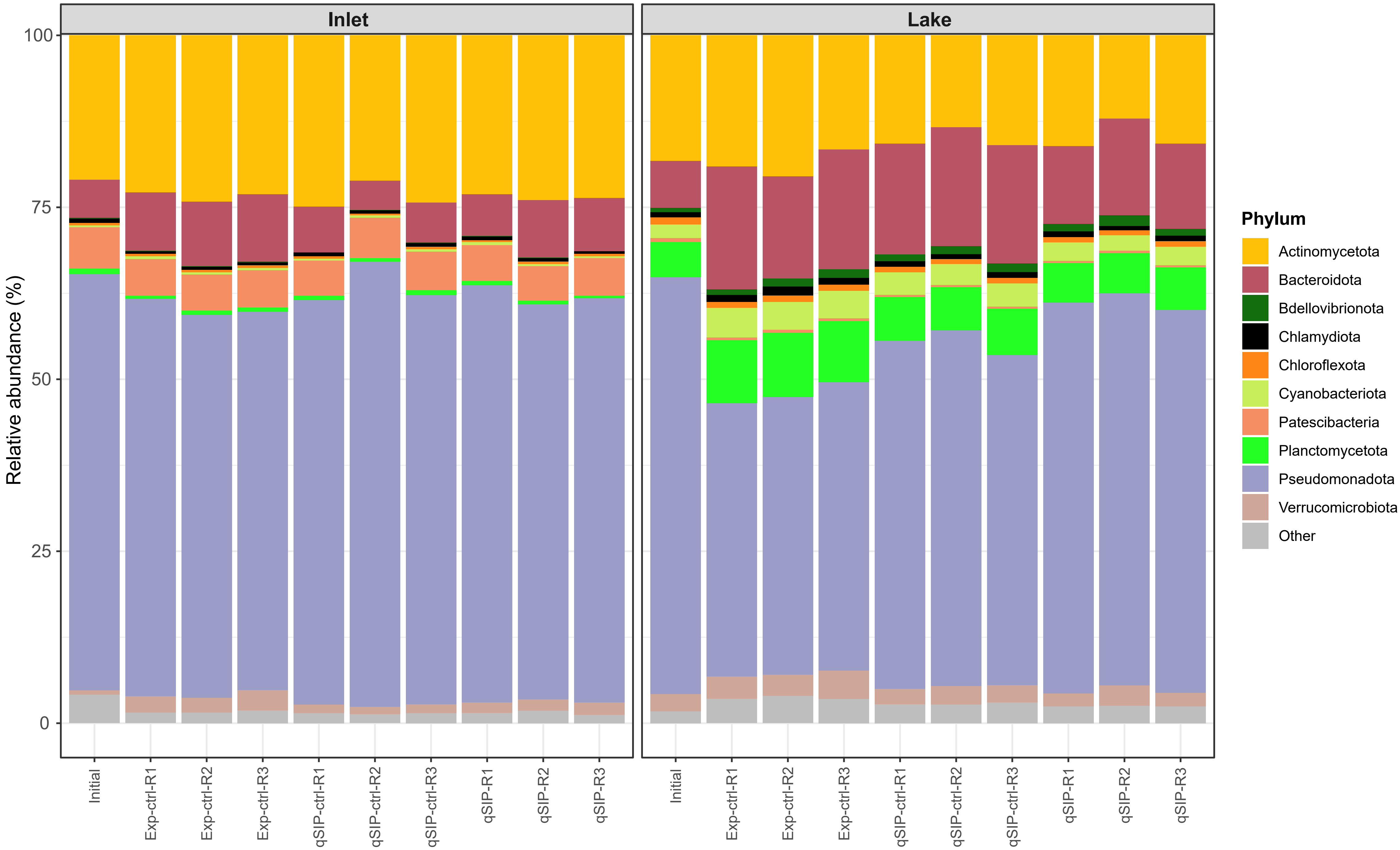


**File S2.** Relative abundance of Operational Taxonomic Units, grouped by Phylum, across all samples sequenced for the 16S rRNA gene. The dataset includes both the initial aquarium communities and the experimental bottle communities from the inlet stream and lake sites. Each bar represents an individual sample, with colors indicating the ten most abundant bacterial Phyla.
