## Supplemental Data 2 for "Protist quantitative stable isotope probing identifies diverse active grazers in natural freshwater communities"

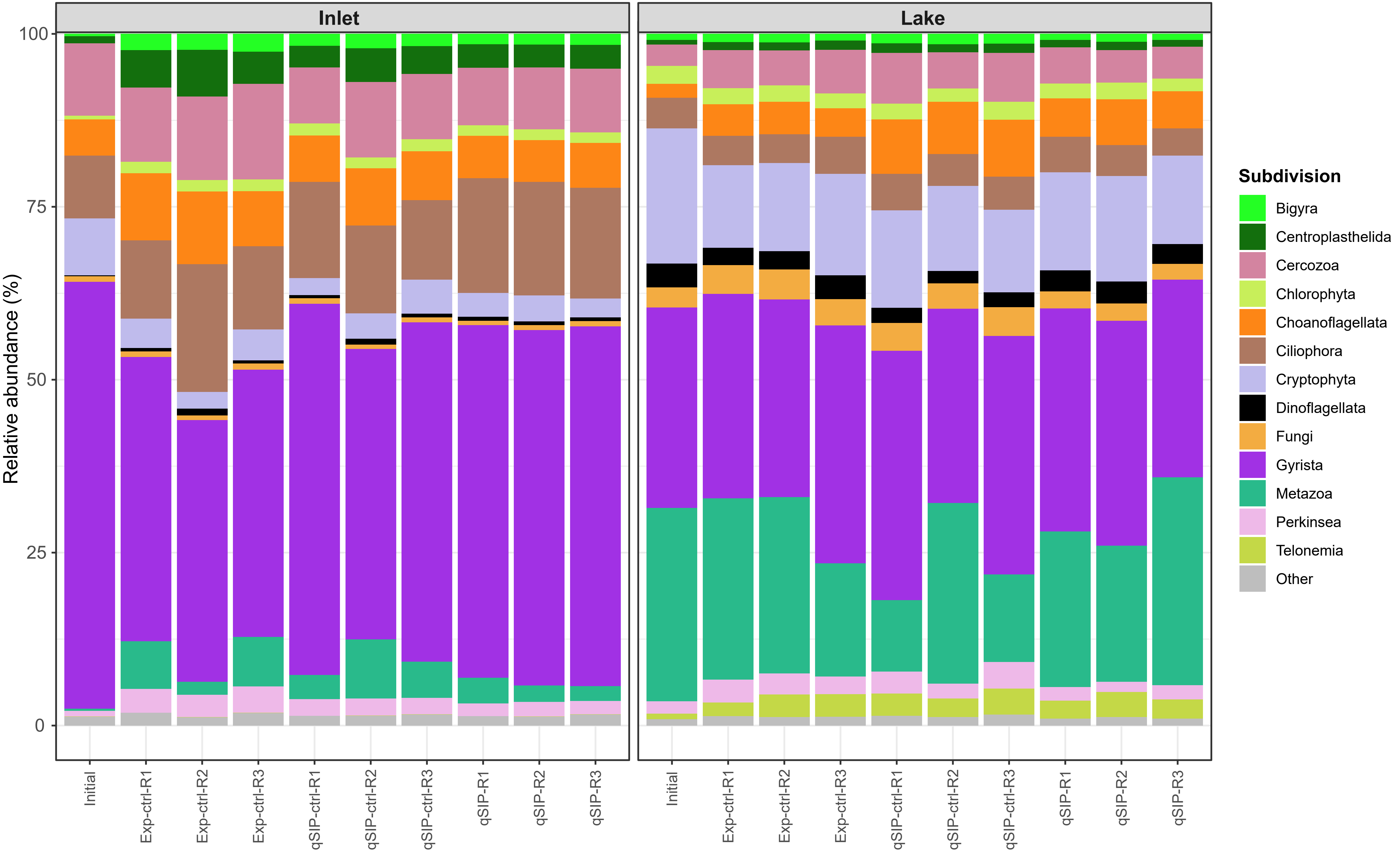


**File S3.** Relative abundance of Operational Taxonomic Units, grouped by Subdivision, across samples sequenced for the 18S rRNA gene. The dataset includes whole community samples (non-fractionated) from both the initial aquarium communities and the experimental bottle communities from the inlet stream and lake sites. Each bar represents an individual sample, with colors indicating the 13 most abundant eukaryotic Subdivisions.
