## Supplementary material for "Protist quantitative stable isotope probing identifies diverse active grazers in natural freshwater communities": sequenced fractions can be found in File S4

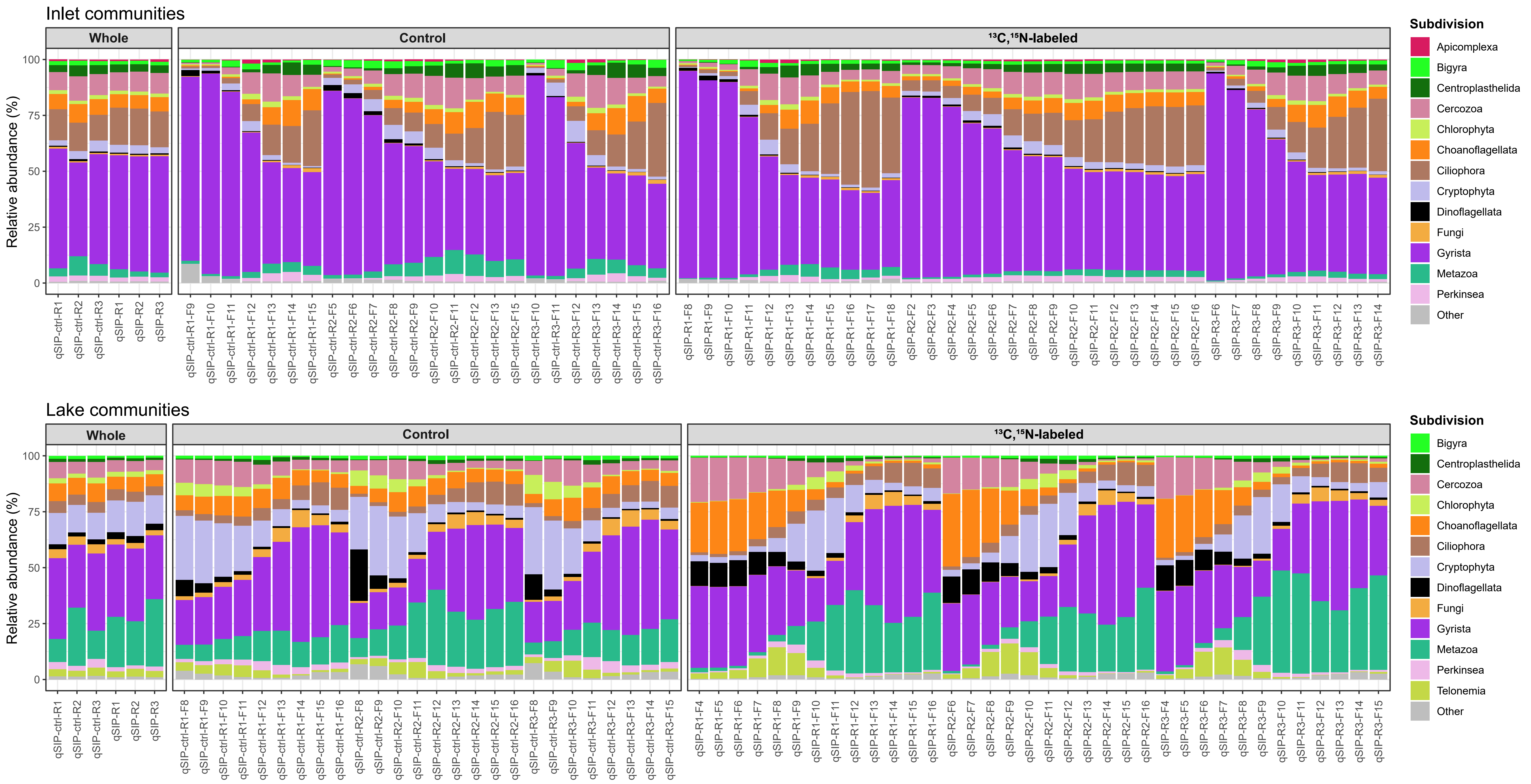


**File S4.** Relative abundance of Operational Taxonomic Units, grouped by Subdivision, across all qSIP-fractions sequenced for the 18S rRNA gene. Fractions (F) from control DNA gradients (unlabeled *Limnohabitans planktonicus* prey) and labeled gradients ($^{13}$C,$^{15}$N- labeled prey) are shown, together with the whole community (non-fractionated) from each bottle incubation for comparison. Each bar represents an individual sample, with colors indicating the 13 most abundant eukaryotic Subdivisions.
