## Supplementary material for "Protist quantitative stable isotope probing identifies diverse active grazers in natural freshwater communities": provided in the Supporting Information, Culture media and prey harvesting

NSY and ISOGRO media were used for the cultivation of *Limnohabitans planktonicus* strain II-D5 (Gram-negative rods, about 0.3–0.4 $\mu$m in diameter and 0.9 $\mu$m long; Kasalický et al. 2010) as prey. After inoculation, cultures were incubated at room temperature (≈ 22 °C) on an Orbital Shaker-Incubator ES-20 (Biosan) at 200 rpm until the desired cell density for prey preparation was reached. ISOGRO medium was filter-sterilized (0.22 $\mu$m, diameter 47 mm; Cytiva Whatman, Nuclepore polycarbonate track-etched membranes), rather than autoclaved, to preserve the integrity of the stable isotope-enriched components, and the same applied to NSY medium for consistency. Media were prepared in 0.5 L autoclaved (121 °C, 15 min) Schott Duran bottles with Duran venting membrane screw caps (0.22 $\mu$m). Detailed compositions of the media are provided in Tables S1–S3.

*L. planktonicus* cells were harvested from cultures using polycarbonate centrifuge tubes (Beckman) in an Avanti J-26 XP centrifuge (Beckman Coulter). Samples were centrifuged at $22 \times 10^3 \times g$ for 12 min at room temperature. After centrifugation, the supernatant was discarded and the pellet was resuspended in 20 mL of the respective culture (NSY or ISOGRO-based). A second centrifugation was performed under the same conditions, followed by a series of washing steps to remove residual medium. Washing steps were performed using inlet or lake water that had been double filter-sterilized (0.22 $\mu$m). After three washes, the final pellets were resuspended in 2 mL of double filter-sterilized inlet or lake water.

Prey abundance in the concentrated stocks was quantified by flow cytometry at $t_0$ (see main text) to calculate the number of prey cells needed to achieve a target abundance equivalent to 20 % of the bacterial community in the experimental bottles. Equal cell numbers were added to the qSIP treatment bottles (with stable isotope-labeled prey) and their respective qSIP controls (with non-labeled prey), ensuring consistent prey abundance within each water type.

### Isotopic composition of *Limnohabitans* cells

A volume of 0.5 L of bacterial culture grown in stable isotope-enriched medium (ISOGRO) was transferred into polycarbonate centrifuge tubes (Beckman) and centrifuged at $22 \times 10^3 \times g$ for 12 min at room temperature using the Avanti J-26 XP centrifuge. The resulting cell pellet was washed twice with phosphate-buffered saline (PBS; pH 7.4, 0.15 M), followed by two additional washes with Milli-Q water. The concentrated cells were collected in a 2 mL Eppendorf tube and immediately frozen at −70 °C. Cells were freeze-dried using a Heto LyoLab 3000 PowerDry freeze dryer. The dried sample was stored in a glass desiccator until delivery to the Tandem Laboratory at Uppsala University for isotopic analysis.

Subsamples of the freeze-dried culture were weighed using a MYA 5 microbalance (Radwag). The relative abundances of stable isotopes ($^{13}C/^{12}C$, $^{15}N/^{14}N$) were determined using an Elementar vario ISOTOPE select elemental analyzer interfaced with an isoprime precisION isotope ratio mass spectrometer.

### Flow cytometry

Prior to analysis, samples and blanks were diluted in PBS as needed. All bacterial samples were pre-filtered through 2 $\mu$m filters (diameter 25 mm; Whatman, Nuclepore). Blanks, used to identify and exclude background noise, consisted of (1) unstained inlet and lake water samples filtered at 0.22 $\mu$m and 2 $\mu$m; (2) 0.22 $\mu$m filtrates stained with SYBR Green I. Lake bacterial samples were diluted fourfold and inlet bacterial samples 16-fold. Flow cytometer settings included a flow rate of 30 $\mu$L min$^{-1}$ and an acquisition until 100,000 events were recorded.

Protist samples were pre-filtered using 70 $\mu$m cell strainers. Blanks included (1) 0.22 $\mu$m, 2 $\mu$m and 70 $\mu$m-filtered unstained inlet and lake water; (2) 0.22 $\mu$m, 2 $\mu$m and 70 $\mu$m-filtered stained samples containing only SYBR Green I or LysoSensor Blue DND-167 (LS). Protist abundance was measured in two size fractions, with samples filtered through 2 $\mu$m and 70 $\mu$m strainers, as performed for the blanks. The 2 $\mu$m fraction was used to correct the 70 $\mu$m fraction for events potentially originating from larger bacteria, so protist abundance corresponds to the 70 $\mu$m fraction minus the 2 $\mu$m events. Feeding ratios were calculated using only the 70 $\mu$m fraction, representing the proportion of LS-positive protists in the large fraction. Lake protist samples were run undiluted, while inlet samples were diluted 16-fold. For these samples, the flow cytometer was operated at a flow rate of 60 $\mu$L min$^{-1}$ with a total acquisition time of 2 min.

### Quantitative PCR

Standard curves for qPCR were generated using a 10-fold serial dilution ($10^8$–$10^1$ gene copies) of PCR-amplified 16S or 18S rRNA genes. The standards were amplified using the same primer sets as the experimental samples and purified from agarose gels (1 %) using the Wizard SV Gel and PCR Clean-Up System (Promega Corporation). The purified amplicons were quantified with a Qubit 3.0 fluorometer (Thermo Fisher Scientific) before preparing the dilution series.

**Table S1.** Composition of the trace element stock solution used in both NSY (unlabeled *L. planktonicus* cells) and ISOGRO (stable isotope-labeled *L. planktonicus* cells) media. The formulation was based on "medium 14" from the German Collection of Microorganisms and Cell Cultures (DSMZ). The solution was filter-sterilized twice through a 0.22 $\mu$m membrane and stored at 4 °C in a polycarbonate bottle until use.

| Component | Weight (g) |
|---|---|
| $Na_2$-EDTA·$2H_20$ | 0.56 |
| $FeSO_4·7H_2O$ | 0.20 |
| $ZnSO_4·7H_2O$ | 0.10 |
| $MnCl_2·4H_2O$ | 0.03 |
| $H_3BO_3$ | 0.30 |
| $CoSO_4·7H_2O$ | 0.24 |
| $CuSO_4·5H_2O$ | 0.015 |
| $NiCl_2·6H_2O$ | 0.02 |
| $Na_2MoO_4·2H_2O$ | 0.03 |
| MQ water | to 1 L |

**Table S2.** Composition of the inorganic solution used in both NSY and ISOGRO media. The formulation follows Hahn et al. (2003). The solution was filter-sterilized twice through a 0.22 $\mu$m membrane and stored at 4 °C until use.

| Component | Amount per L of water |
|---|---|
| $MgSO_4 \cdot 7H_2O$ | 75 mg |
| $Ca(NO_3)_2 \cdot 4H_2O$ | 43 mg |
| $NaHCO_3$ | 16 mg |
| KCl | 5 mg |
| $K_2HPO_4 \cdot 3H_2O$ | 3.7 mg |
| Trace element solution (see Table S1) | 100 $\mu$L |
| Milli-Q water | to 1 L |

**Table S3.** Composition of the NSY medium used for cultivation of *L. planktonicus* prey (without stable isotopes) based on Hahn et al. (2003). The medium had a measured pH of 7.2, measured using a VWR pHenomenal pH 1100L/1100LB pH/mV/°C bench meter at ambient temperature, and no adjustment was required. The medium was filter-sterilized twice through a 0.22 $\mu$m membrane and stored at 4 °C in autoclaved glass bottles until use.

| Component | Weight (g) |
|---|---|
| Nutrient broth | 1 |
| Peptone from soybean | 1 |
| Yeast extract | 1 |
| Inorganic solution (see Table S2) | to 1 L |

**Table S4.** qPCR cycling conditions for 18S rRNA gene copy number quantification.

| Step | Temperature (ºC) | Duration | Cycles |
|---|---|---|---|
| Initial denaturation | 98 | 3 min | 1 |
| Denaturation | 98 | 15 sec | 40 |
| Annealing | 52 | 40 sec | |
| Extension | 72 | 40 sec | 1 |

**Table S5.** qPCR cycling conditions for 16S rRNA gene copy number quantification to assess the extent of labeling of *Limnohabitans planktonicus* cells grown in the stable isotope-enriched medium (ISOGRO).

| Step | Temperature (ºC) | Duration | Cycles |
|---|---|---|---|
| Initial denaturation | 95 | 3 min | 1 |
| Denaturation | 95 | 10 sec | 40 |
| Annealing/Extension | 55 | 30 sec | |

**Table S6.** Primers used for the amplicon sequencing libraries. The 16S rRNA V3-V4 primers included phasing variants and were used for library preparation by the National Genomics Infrastructure (NGI; Stockholm, Sweden). The 18S rRNA V4 primers included four random nucleotides (NNNN) to the 5' end of the forward primer.

| Gene/region | Primer | Sequence (5' to 3') | Comment | |
|---|---|---|---|---|
| 16S rRNA/ V3-V4 | 341F | CCTACGGGNGGCWGCAG | Forward primer | phasing 0 |
| | | **t**CCTACGGGNGGCWGCAG | | phasing 1 |
| | | **gt**CCTACGGGNGGCWGCAG | | phasing 2 |
| | | **agt**CCTACGGGNGGCWGCAG | | phasing 3 |
| | | **gagt**CCTACGGGNGGCWGCAG | | phasing 4 |
| | | **agagt**CCTACGGGNGGCWGCAG | | phasing 5 |
| | | **tagagt**CCTACGGGNGGCWGCAG | | phasing 6 |
| | | **ctagagt**CCTACGGGNGGCWGCAG | | phasing 7 |
| | 805R | GACTACHVGGGTATCTAATCC | Reverse primer | phasing 0 |
| | | **g**GACTACHVGGGTATCTAATCC | | phasing 1 |
| | | **tg**GACTACHVGGGTATCTAATCC | | phasing 2 |
| | | **ctg**GACTACHVGGGTATCTAATCC | | phasing 3 |
| | | **actg**GACTACHVGGGTATCTAATCC | | phasing 4 |
| | | **tactg**GACTACHVGGGTATCTAATCC | | phasing 5 |
| | | **ctactg**GACTACHVGGGTATCTAATCC | | phasing 6 |
| | | **actactg**GACTACHVGGGTATCTAATCC | | phasing 7 |
| 18S rRNA/ V4 | V4F | **NNNN**CCAGCASCYGCGGTAATTCC | Forward primer with four Ns | |
| | V4RB | ACTTTCGTTCTTGATYRR | Reverse primer | |

**Table S7.** PCR1 cycling conditions for the 18S rRNA gene amplicon library, adapted from Latz et al. (2022). PCR reactions were performed using the KAPA HiFi HotStart ReadyMix PCR Kit (Kapa Biosystems).

| Step | Temperature (ºC) | Duration | Cycles |
|---|---|---|---|
| Initial denaturation | 95 | 3 min | 1 |
| Denaturation | 98 | 20 s | 24 |
| Annealing | 52 | 15 s | |
| Extension | 72 | 15 s | |
| Final extension | 72 | 2 min | 1 |

**Table S8.** Measurements of water temperature, dissolved oxygen concentration and oxygen saturation in situ and in the aquaria used for water storage prior to the grazing experiment.

| Site | Depth (m) | Temperature (ºC) | Dissolved oxygen (mg $L^{-1}$) | Oxygen saturation (%) |
|---|---|---|---|---|
| | | ***In situ*** | | |
| **Inlet** | 0-0.3 | 17.7 | 2-5 | NA* |
| **Lake** | 0 | 19.0 | 8.81 | 93.8 |
| | 0.5 | 18.4 | 8.92 | 93.9 |
| | 1 | 18.1 | 8.91 | 93.3 |
| | 1.5 | 17.8 | 8.76 | 91.1 |
| | 2 | 17.6 | 8.63 | 89.5 |
| | 2.5 | 17.5 | 8.62 | 89.0 |
| | 3 | 17.4 | 8.59 | 88.5 |
| | 3.5 | 17.3 | 8.54 | 87.9 |
| | 4 | 17.0 | 8.19 | 83.7 |
| | | **Aquaria** | | |
| **Inlet** | 0 | 16.5 | 7.43 | 74 |
| **Lake** | 0 | 17 | 9.56 | 97.4 |

***** Unstable measurement

**Table S9.** Concentrations of total organic carbon (TOC), total nitrogen (TN) and total phosphorus (TP) at the beginning of the experiment ($t_0$, single measurement) and after 36 h of incubation (mean ± standard deviation of three replicates). The standard deviation of TP measurements in the inlet stream was relatively high, which may be attributed to the elevated turbidity of the inlet water.

| Site | Incubation | TOC (mg $L^{-1}$) | TN (mg $L^{-1}$) | TP ($\mu$g $L^{-1}$) |
|---|---|---|---|---|
| **Inlet** | $T_0$ (*n*=1) | 26 | 1.2 | 160 |
| | Experiment control (*n*=3) | 23±0.2 | 0.93±0.03 | 160±60 |
| | qSIP control (*n*=3) | 22±0.1 | 0.88±0.02 | 200±20 |
| | qSIP (*n*=3) | 25±6 | 1.1±0.5 | 150±30 |
| **Lake** | $T_0$ (*n*=1) | 15 | 0.43 | 14 |
| | Experiment control (*n*=3) | 16±0.1 | 0.43±0.01 | 12±1 |
| | qSIP control (*n*=3) | 15±0.1 | 0.44±0.01 | 12±0.3 |
| | qSIP (*n*=3) | 15±0.2 | 0.43±0.01 | 15±0.9 |

**Table S10.** Analysis of Bray-Curtis dissimilarities among whole (non-fractionated) 16S rRNA gene communities (*n* = 20), and whole and qSIP-fractionated 18S rRNA gene communities (*n* = 141). Differences in dispersion were assessed using the *betadisper* function, followed by a permutation-based test. PERMANOVAs were conducted using the *adonis2* function (both from the vegan package in R). The NMDS stress value is also provided. "Site" refers to inlet stream or lake water samples sequenced.

| Group | Number of OTUs | Factor | | **Dispersion** (*betadisper*) | | **PERMANOVA** (*adonis2*) | | | NMDS |
|---|---|---|---|---|---|---|---|---|---|
| | | | df | F value | *p* value | F value | $R^2$ | *p* value | **stress** |
| Bacterial communities | 10,705 | Site | 1,18 | 0.69 | 0.448 | 145.13 | 0.890 | **<0.001** | 0.00008 |
| Eukaryotic communities | 2,560 | Site | 1,139 | 4.53 | **0.028** | 154.87 | 0.527 | **<0.001** | 0.049 |

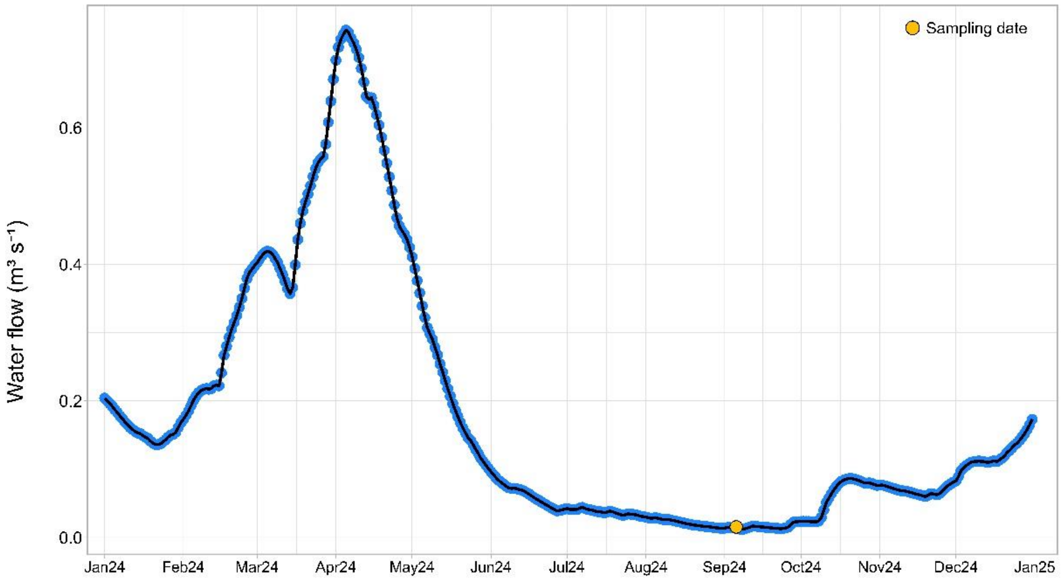


**Figure S1.** Model-estimated daily total water flow in 2024 at the outlet of lake Siggeforasjön, with the sampling date indicated.

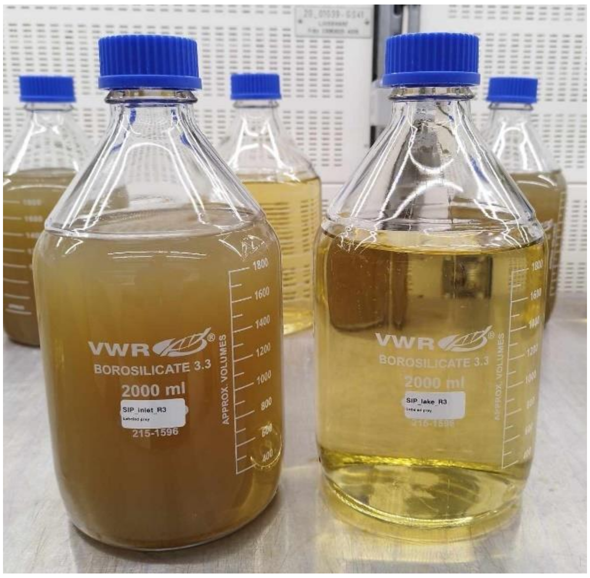


**Figure S2.** Two of the experimental bottles containing inlet stream water (left) and lake pelagic water (right). The higher haziness of the inlet sample is apparent.

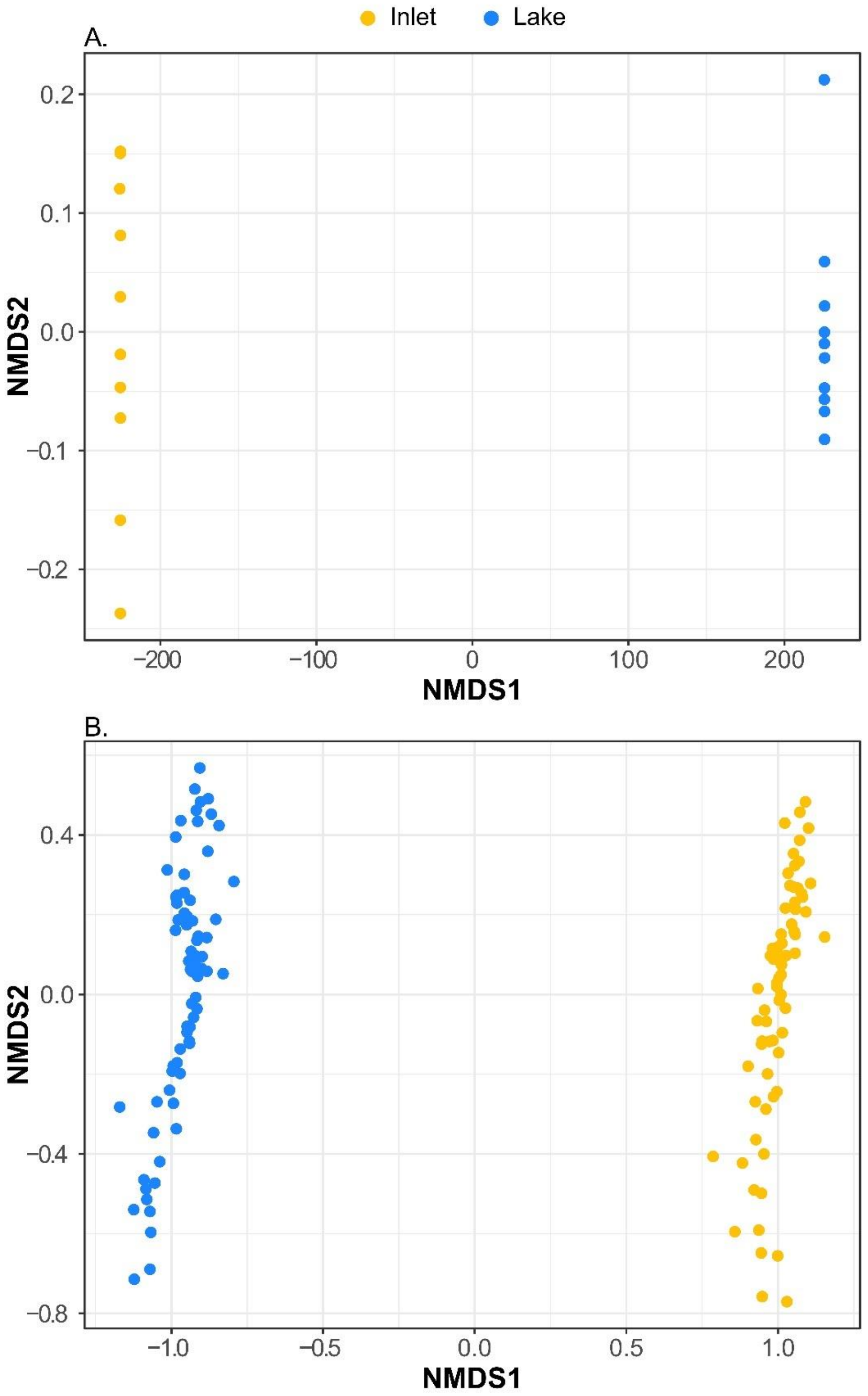


**Figure S3.** Non-metric multidimensional scaling (NMDS) plots, showing (A) bacterial (16S rRNA; top) and eukaryotic (18S rRNA; bottom) communities. Each symbol represents an individual sample (*n* = 20 for bacteria; *n* = 141 for eukaryotes) with colors indicating the sampling site. For eukaryotes, both whole (non-fractionated) and qSIP-fractionated samples are included in the NMDS.

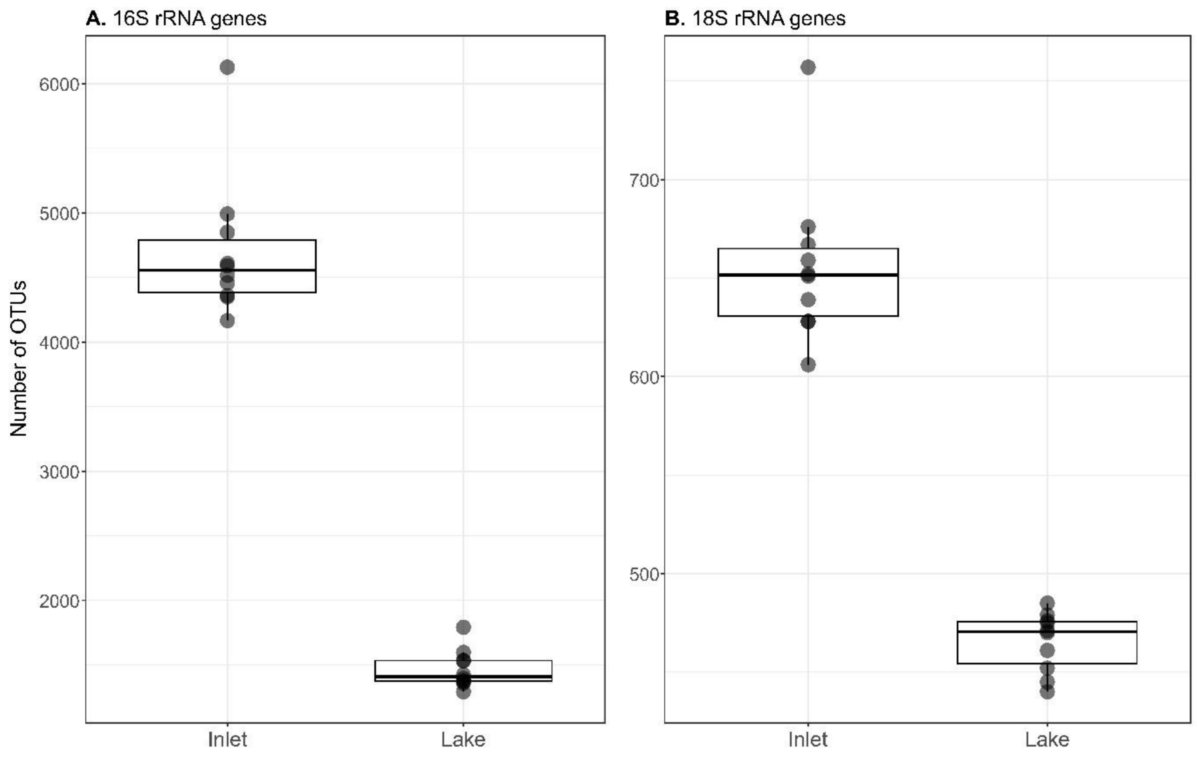


**Figure S4.** Observed species richness as the number of operational taxonomic units (OTUs) in (A) bacterial (*n* = 10 per site) and (B) eukaryotic (*n* = 10 per site) whole communities. For the 16S rRNA dataset, reads were rarefied to 508,072 reads per sample, while for the 18S rRNA reads were rarefied to 377,024 reads per sample.

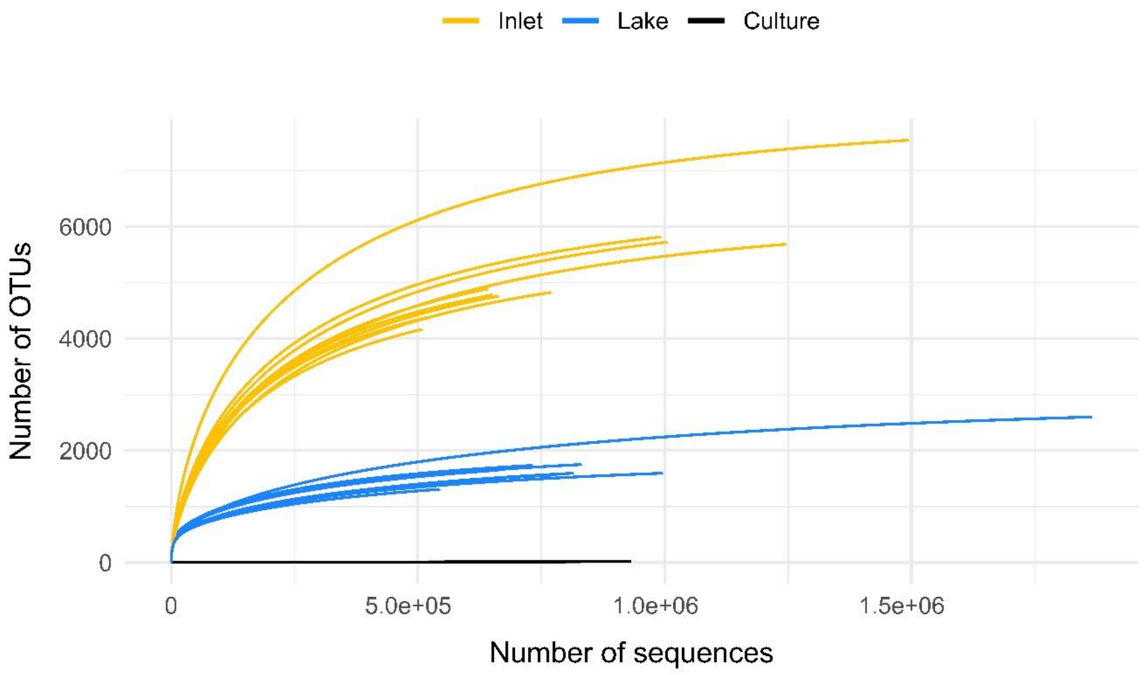


**Figure S5.** Rarefaction curves of bacterial communities (16S rRNA gene sequences). Curves show sequencing depth across samples with inlet, lake and culture samples distinguished by color.

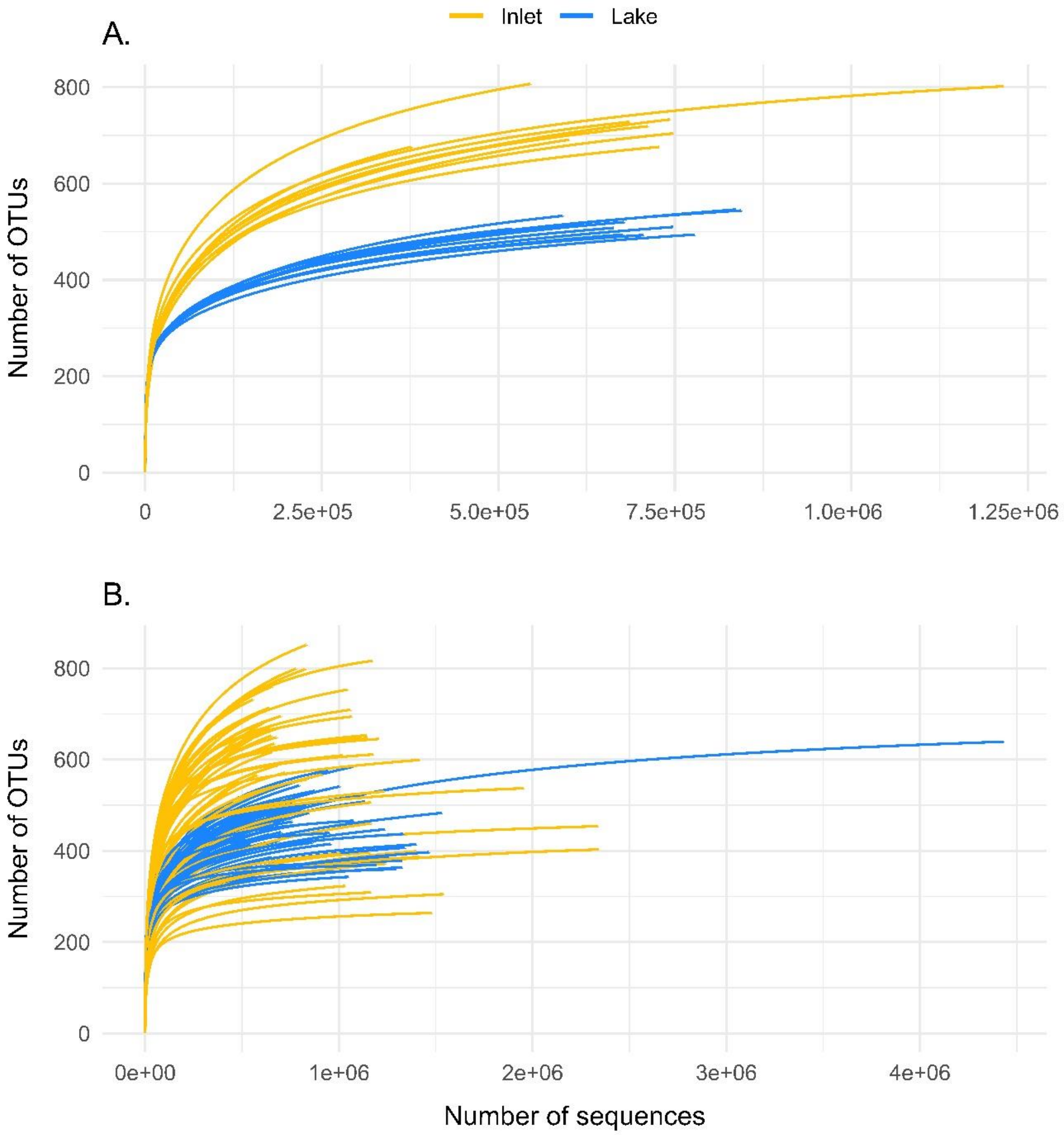


**Figure S6.** Rarefaction curves of eukaryotic communities (18S rRNA gene sequences). Curves show sequencing depth across samples for (A) whole (non-fractionated) eukaryotic communities and (B) samples fractionated for quantitative stable-isotope probing (qSIP). Inlet and lake samples are distinguished by color.

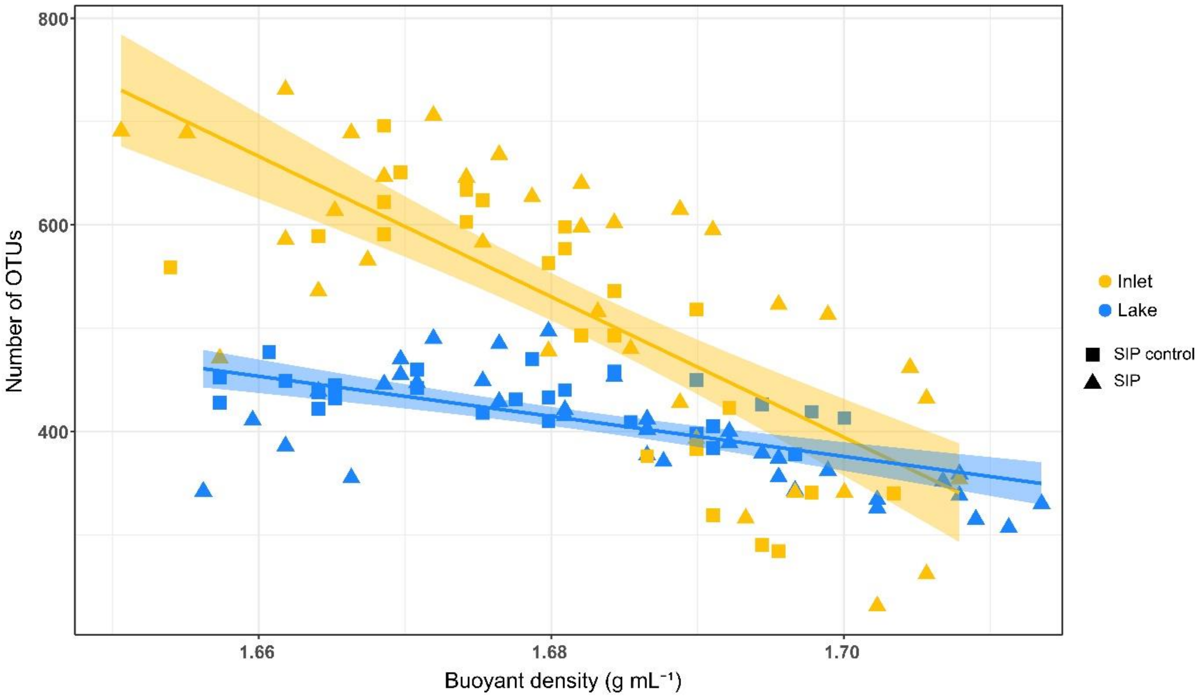


**Figure S7.** Relationship between eukaryotic OTU richness from qSIP-fractionated samples and buoyant density. Lines represent linear regressions, colors the site and point shapes the incubations with unlabeled prey ("SIP control") or $^{13}C$, $^{15}N$-prey ("SIP"). The 18S rRNA reads were rarefied to 377,024 reads per sample.

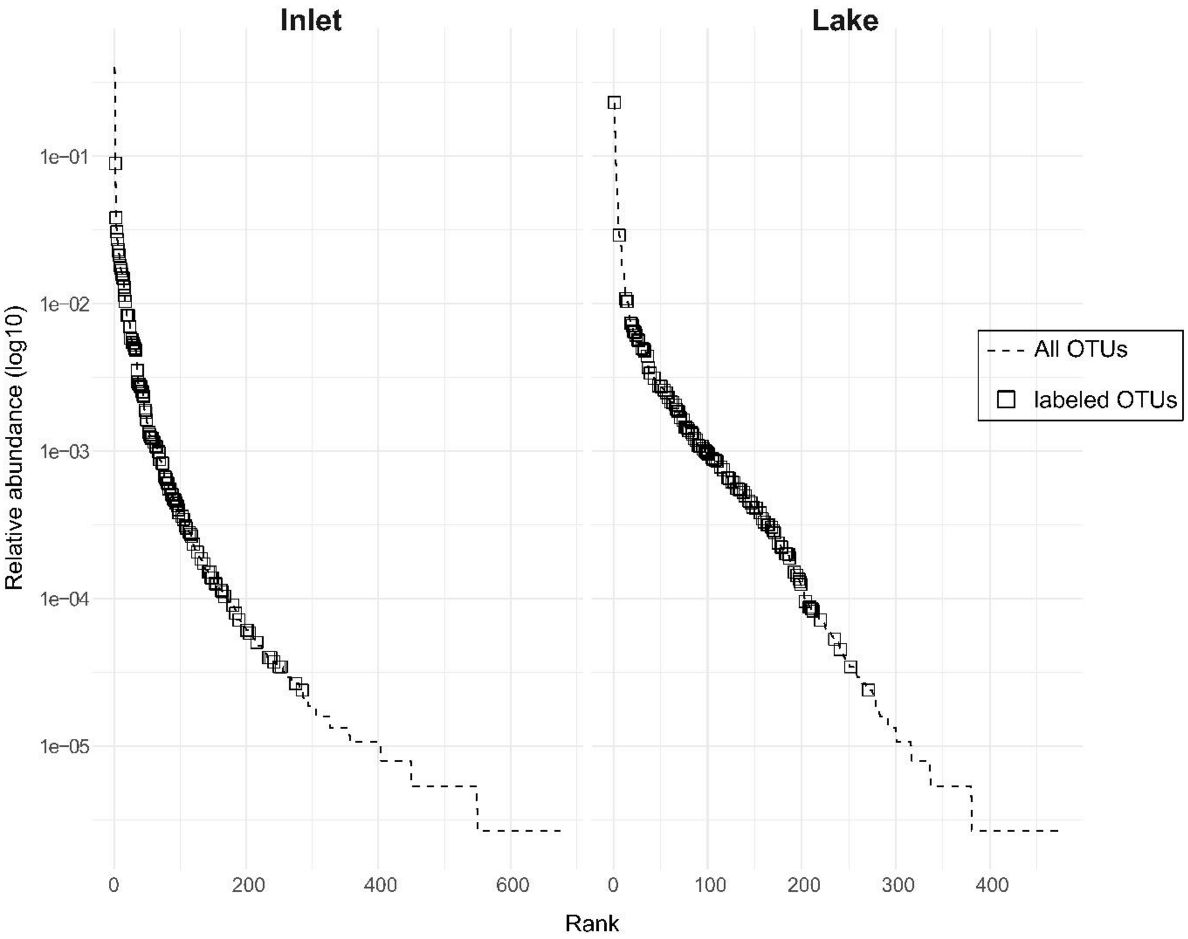


**Figure S8.** Rank abundance plots for the inlet stream (left) and lake (right) 18S rRNA gene communities sampled from the aquaria at $t_0$. Dashed lines represent rank abundance curves based on rarefied data to 377,024 reads, resulting in 676 OTUs for the inlet and 475 OTUs for the lake sample. Each OTU was assigned a unique rank. Labeled OTUs identified through qSIP are shown as squares ($n$ = 108 for the inlet and $n$ = 107 for the lake).
